# Calcium Dependent Conformational Changes in Human Transglutaminase 2 and its Implications in Celiac Disease

**DOI:** 10.1101/2025.06.04.657948

**Authors:** Srimari Srikanth, Pavinap Priyaa Karthikeyan, Mezya Sezen, Dharineesh K Sridhar, Karthikeya Doppalapudi, Taner Z. Sen, Venkatasubramanian Ulaganathan, Ragothaman M. Yennamalli

**Affiliations:** Department of Bioinformatics, School of Chemical & Biotechnology, SASTRA Deemed to be University, Tirumalaisamudram, Thanjavur, Tamil Nadu 613401, India; Molecular Motors Lab, Department of Biotechnology, School of Chemical & Biotechnology, SASTRA Deemed to be University, Tirumalaisamudram, Thanjavur, Tamil Nadu 613401, India; U. S. Department of Agriculture, Agricultural Research Service, Crop Improvement and Genetics Research Unit, 800 Buchanan Street, Albany, California 94710, United States of America; Department of Bioengineering, University of California, Berkeley, CA, 94720, United States of America

**Keywords:** Molecular Dynamics simulation, Coarse-Grained Dynamics, Celiac Disease, Calcium-dependency, Transglutaminase 2

## Abstract

Transglutaminase 2 (TG2) serves as a modifiable transamidating acyltransferase that precipitates calcium-induced protein alterations. The enzyme plays a crucial role in the cell and disease states, such as tissue repair, calcium signal transduction, celiac disease, and cancer. It is implicated in protein crosslinking and has been found in high concentrations in the small intestines of those with celiac disease. The function of TG2 hinges upon calcium ions binding to particular sites on the enzyme. In this study, we delve into the contribution of calcium-responsive transglutaminase 2 (TG2) in celiac disease, utilizing both molecular dynamics simulations and coarse-grained models, and investigate the impact of non-synonymous single nucleotide polymorphisms (nsSNPs) on TG2. Molecular dynamics reveal prominent conformational differences between the open and closed conformations. In the coarse-grained model, key residues are found adjacent to the active site in the open conformation, while in the closed conformation, key residues are distant from the active site. We further explore the functional impact of non-synonymous single nucleotide polymorphisms (nsSNPs) in TG2 using both sequence-based and structure-based computational tools. Through a consensus approach, we identify ten nsSNPs that are predicted to destabilize TG2 or alter its structural flexibility, with mutations such as R48H, E186Q, C277S, and E549G likely to influence active site accessibility and calcium coordination. The findings from this research enhances our understanding of the molecular processes underpinning celiac disease and helps facilitate innovative treatment approaches that target calcium-responsive TG2.

## Introduction

Transglutaminase 2 (TG2), a calcium (Ca^2+^) reliant enzyme, belongs to the transglutaminase family and has a mass of 78 kDa (Klock et al., 2012). It is prevalently present in both intracellular and extracellular compartments across various organs, including the liver, intestine, skin, and brain (Lorand et al., 2003). The enzyme is implicated in a wide spectrum of physiological functions, such as wound healing, cell differentiation (Balajthy. et al., 2006), and apoptosis (Klöck et al., 2012) and encoded by the TGM2 gene, located on the 20^th^ human chromosome.

TG2 facilitates protein cross-linking and deamination through the post-translational amendment of glutamine residues located on the protein substrate (Klock et al., 2012). Within TG2, a thiol group of cysteine residue initiates the cross-linking process and interacts with a surface glutamine residue, discharging ammonia and establishing a thioester intermediate. Consequently, an isopeptide bond between the two substrates emerges when a second substrate’s surface amine attacks the thioester intermediate. The hydrolysis of thioester intermediate can potentially transform the glutamine residue into glutamic acid (Folk et al., 1987). When TG2 is dysregulated, it can trigger several conditions, including celiac disease (Dieterich et al., 1997), Alzheimer’s disease (Wang et al., 2008), and cancer (Mangala and Mehta, 2005).

TG2 contains four domains (Figure 1) namely the β-sandwich domain at the N-terminal, the catalytic (core) domain, and two β-barrels (also known as β1 and β2 domains) at the C-terminal. TG2 has two conformation states, an extended conformation (open conformation) (Figure 1A), and the compact conformation (closed conformation) (Figure 1B) (Pinkas *et al*., 2007). The superposition of the open and closed conformations of TG2 (Figure 1C) indicates that the domains (N-terminal and core domain) of both conformations are structurally aligned. However, notable large conformational differences are observed between the two C-terminal β-barrels domains. TG2 can exist in multiple conformations, including the open and closed states, depending on its binding to various ligands, such as calcium ions and substrate proteins (Liu et al., 2002). In the open conformation, gluten peptide is bound to the active site whereas in the closed conformation guanine diphosphate (GDP) or adenine triphosphate (ATP) is bound to the active site.

**Figure 1:**
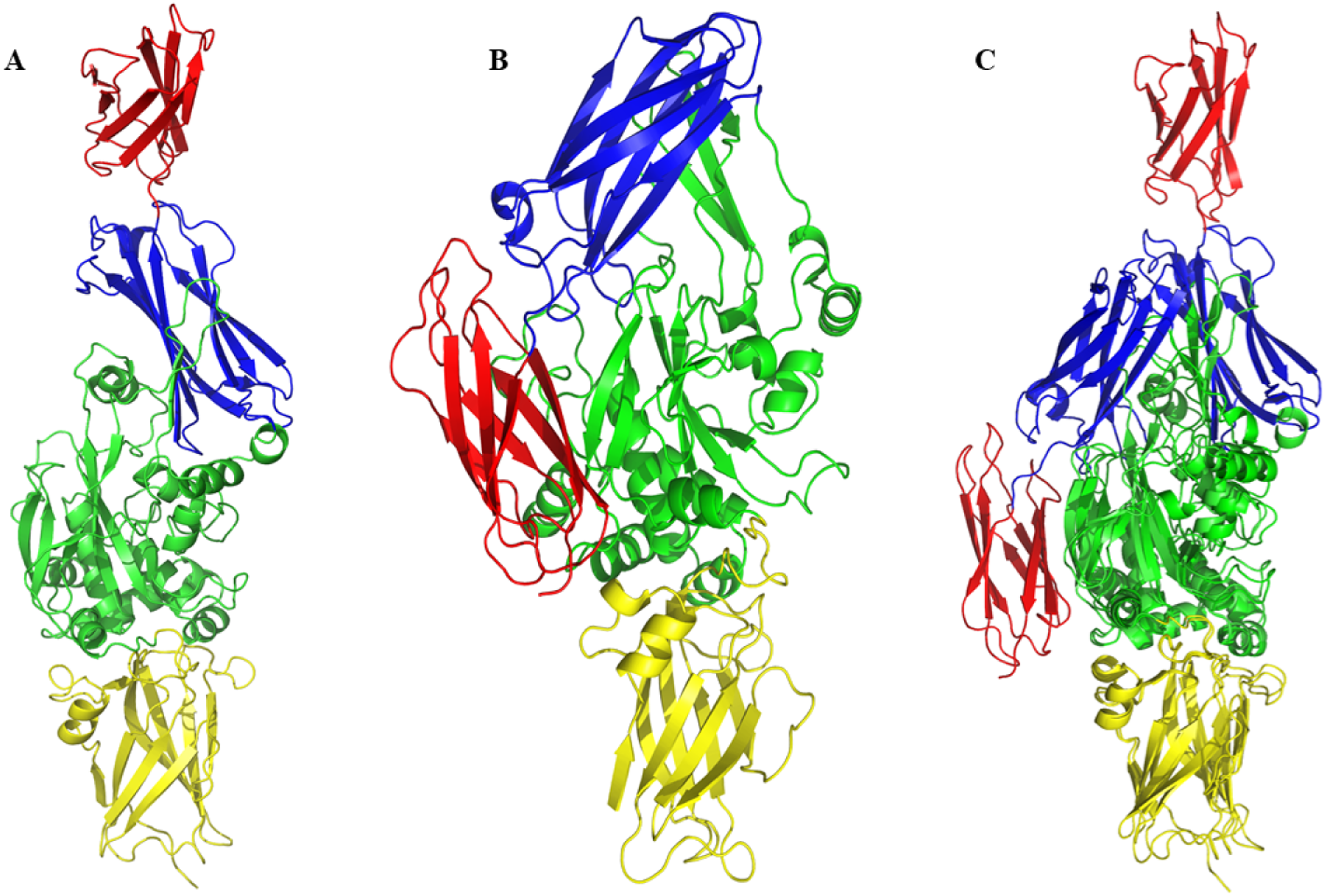
Conformation of TG2 structures available in PDB. (A) The open conformation of TG2, (B) its closed conformation and (C) the superposition of both conformations. The N-terminal β-sandwich domain is shown in yellow, the catalytic domain in green, β1 in blue, and β2 domain in red. This representation was generated using PyMOL (Schrodinger and DeLano 2017)

The specific mechanisms of interactions between Ca^2+^ and TG2 still elude a comprehensive understanding (Jeong et al., 2020). TG2 is understood to assume a yet undefined native conformation, as outlined by (Staffler et al., 2020). The precise pathway leading to TG2’s activation has not been completely characterized and the characteristics of the intermediate TG2 conformations also remain unknown (Király et al., 2009).

Celiac disease is an autoimmune disorder caused by the gluten, specifically affecting the small intestine. The immune system recognizes gluten as a foreign particle and mounts an immune response against it. Consequently, this leads to inflammation and damage to the finger-like projection called villi in the small intestine, which is essential for the nutrient absorption (Rubio-Tapia *et al*., 2013). Examining findings from anti-tissue transglutaminase tests unveils a global occurrence rate of celiac disease stands at approximately 1.4%, affecting an estimated 275,818 individuals (Singh *et al*., 2018). In India, celiac disease has garnered recent recognition and awareness. One contributing factor to the underdiagnosis of celiac disease in India is the diverse range of dietary patterns across the country, affecting one percent of the North Indian population. Symptoms of celiac disease can vary and often overlap with those of other digestive disorders, making it challenging to diagnose without proper testing. (Kumar *et al*., 2017)

Abdominal pain, bloating, diarrhea, weight loss, exhaustion, and vitamin and mineral deficiencies are among the many symptoms of celiac disease. People who have celiac disease often develop extra-gastrointestinal symptoms such as skin rashes, pain in the joints, and neurological problems (Holtmeier and Caspary, 2006).

The open conformation of TG2 is understood to play an important role in celiac disease. The gluten peptide enters a cell and binds to the TG2 enzyme. The TG2 gets activated, and the conformations change from the closed conformation to the open conformation (Kim and Park, 2020). This process causes the deamidation of gluten, which acts as a foreign particle. These glutens cause the activation of anti-transglutaminase antibodies and affect the small intestine (Caja *et al*., 2011).

The following findings support the above idea, (i) TG2 prefers for specific gluten peptides that are known to provoke immune reactions, (ii) deamidation process catalyzed by TG2 greatly increases the recognition of these gluten peptides by HLA-DQ2 or HLA-DQ8, both of which are closely associated with the disease (iii) When gluten-responsive T cells from the celiac intestine are activated, they produce large quantities of interferon-γ, an essential signaling molecule. Also, interferon-γ triggers extracellular TG2 activation through a TRX-mediated mechanism, potentially creating a self-amplifying loop that exacerbates gluten-induced inflammation (Rauhavirta et al., 2019). Further, continuous exposure of celiac patients to dietary gluten consistently leads to the production of autoantibodies against TG2 (Kloeck, C. 2014). While the mechanisms behind some of these phenomena are well-understood, others remain less clear. Therefore, in depth investigation into these TG2 related process is needed to know the role of transglutaminase activity in the pathogenesis of celiac disease but also reveal new insights into the fundamental biological functions of TG2 in the human body.

This investigation seamlessly transitions into an examination of non-synonymous single nucleotide polymorphisms (nsSNPs) within TG2, showcasing their disruptive potential and how alterations in TG2’s structure and function due to nsSNPs can impact its interactions with gluten peptides, potentially influencing the development or severity of celiac disease (Figure 15). Non-synonymous single nucleotide polymorphisms (nsSNPs) within TG2 instigates alterations in the protein sequence (Thangaraju et al., 2017). The disruptive potential of such extends to the structure, function, and regulation of TG2, subsequently influencing its physiological operations (Kiraly et al., 2011). The identification and characterization of nsSNPs in the TG2 gene have gained significant attention, particularly concerning their potential implications for celiac disease. Any alterations in the structure or function of TG2 due to nsSNPs have potential to impact its ability to interact with gluten peptides, potentially contributing to the development or severity of celiac disease (Dieterich *et al.,* 1997). This involves exploration of how these variations affect the enzymatic activity, protein-protein interactions, cellular localization, and overall function of TG2 (Zemskov *et al.,* 2006). Understanding the impact of TG2 nsSNPs is significant in elucidating the molecular mechanisms underlying these diseases, identifying individuals at higher risk, and potentially developing targeted therapeutic approaches. Through an in-depth investigation into the functional consequences of TG2 nsSNPs, these aim to gain insights into the role of TG2 variants in disease susceptibility, prognosis, and treatment response. This, in turn, contributes to advancement of personalized medicine approaches tailored to individual patients.

Molecular dynamics (MD) is a computational technique used to explore protein dynamics at the atomic level (Hollingsworth and Dror, 2018). Performing these simulations demands substantial computational resources due to the necessity for energy calculations at the femtosecond scale, i.e., 10^-15^ seconds, to reach a biologically significant timescale exceeding 10^-6^ seconds (Hildebrand et al., 2019). These simulations rely on classical physics principles, specifically Newton’s laws of motion, applied to each atom individually. Forces are determined by the original structural arrangement, positioning, and velocities of each atom (Berendsen et al., 1995). These forces are quantified using mathematical formulas that govern atom interactions, covering both bonded and non-bonded interactions. By numerically integrating these equations, MD simulations can trace the position, velocity, and energy of individual particles over time (Karplus and McCammon, 2002). Force fields include parameters describing covalent bond extension, angle distortion, torsion angles, and non-bonded interactions such as van der Waals forces and electrostatic interactions. These interactions and the strategic handling of solvent effects enable the simulation of intricate biological systems (Wu et al., 2022). MD simulations offer insights into protein dynamics, folding mechanisms, conformational transitions, and ligand binding processes. Additionally, MD simulations can evaluate the impacts of temperature, pH, mutations, and other factors on the stability and architecture of proteins (Patodia, 2014).

Coarse-grained (CG) dynamics is a computer modelling method that simplifies the representation of complex systems by defining a residue as a single entity rather than focusing on individual atoms (Riniker et al., 2012). In contrast to MD simulations, the CG method excels in modelling the longer timescale dynamics of macromolecules in a short time by sacrificing atomistic-level details (Bahar *et al*., 1997). This approach holds the potential to describe slow collective motions and long-range interactions, which are difficult to analyze in atomistic simulations. The simplified representation of the system allows for the exploration of processes that occur place over longer periods, such as those that last for milliseconds (Doruker *et al.,* 2000).

CG dynamics in proteins that show large conformational domain movements, and in proteins that bind to metals. Various CG methods available are available to perform dynamics., and one frequently employed for coarse-grained modelling is the Elastic Network Model (ENM). In the ENM method, each residue is considered a node (C^α^), and nodes are inter-connected by springs (Bahar *et al*., 1997). Another method, Anisotropic Network Model (ANM), serves to determine the directionality of motion (Atilgan *et al.,* 2001). Meanwhile, Gaussian Network Model (GNM) methods calculate a mean square fluctuation value (Kundu *et al*., 2002), which is comparable to the experimental B-factor or temperature factors also known as Debye-Waller factor. ENM excels in capturing slow functional motions and understanding the global behavior of a protein. It provided valuable insights into the regions crucial for functionality. Complementarily, Molecular Dynamics (MD) can offer the same information but at a more detailed, atomic level. Together, these methods contribute to a comprehensive understanding of the dynamic behavior of proteins.

Our models depict domain-level conformational changes in TG2 with the impact of Ca2+ ions. In a comprehensive analysis of TG2 nsSNPs, we identified 10 variants using sequence and structure-based tools, highlighting their destabilizing or deleterious nature. This work establishes a foundation for future computational and experimental studies for a deeper understanding of TG2 function and structural dynamics.

## Materials and Methods

We established a computational workflow to study human TG2 (Figure 13), utilizing the protein information sourced from the Protein DataBank (PDB) database. We modelled the protein structure in SWISS-MODEL (Waterhouse *et al.,* 2018),and subsequently developed four setups for molecular dynamics (MD) simulations. We used crystal structures to build a coarse-grained (CG) model to investigate the behavior of TG2 and evaluate the impact of mutations on protein structures, allowing for a thorough exploration of TG2 dynamics and the effect of variations on dynamics.

### Structures Selection

We conducted a search for human TG2 in the PDB database by using the query: "*tgm2*" and selecting *Homo sapiens* as source organism. We found nine crystal structures of TG2 in the PDB database. Among these structures, five are in closed conformation with GDP or ADP bound (PDB id: 1KV3, 3LY6, 4PYG, 6KZB, and 6A8P), and three are in open conformation with N-[(2R)]. -2-{[(benzyloxy)carbonyl]amino} -7-ethoxy-7-oxoheptanoyl] -L-valyl-L-prolyl-L-leucine (PDB id: 3S3S), N-[(benzyloxy)carbonyl] -6-diazonio-5-oxo-L-norleucyl-L-valyl-L-prolyl-L-leucine (PDB id: 3S3J), and N-2-[(2S)-2-[(benzyloxy)carbonyl]amino -7-ethoxy-7-oxoheptanoyl] -L-glutaminyl-L-prolyl-L-leucine (PDB id: 3S3P) peptides bound, and one is in an open conformation with Ca^2+^ (PDB id: 2Q3Z).

### Modelling of structures

We found many missing residues in all structures (Supplementary Tables 1 and 2). Given that structures with missing residues are not recommended for detailed fine-grained analysis, we used the monomer structure and filled the missing residues computationally using SWISS MODEL (Waterhouse et al., 2018).

The superimpositions of both modelled and crystal structures are shown in Figure 9. The root-mean-square-deviation (RMSD) between the modelled TG2 structure in the open conformation and its corresponding crystal structure (PDB id: 2Q3Z) was 0.058 Å. Similarly, the RMSD between the modelled TG2 structure in the closed conformation and its corresponding crystal structure (PDB id: 6KZB) was 0.381Å. The open and closed conformations in complex with and without two Ca^2+^ ions were built using PyMOL. Specifically, the coordinates of Ca^2+^ ion were transferred from 6KZB crystal structure to the modelled proteins. These four setups were used for subsequent MD simulations.

### Coarse Grained Dynamics

We used ProDy (protein dynamics) for coarse-grained simulation (Zhang *et al*., 2021). Specifically, the elastic network model (ENM) was implemented for coarse-grain analysis, incorporating two network model methods: the Gaussian Network Model (GNM) and the Anisotropic Network Model (ANM). GNM was utilized to monitor global dynamics, representing the protein structures as a C^α^ network connected by springs within a distance of 7 Å for GNM and 13 Å for ANM, standard parameters for these models (Atilgan et al., 2001).

The fluctuations are assumed to follow a Gaussian distribution, with each residue depicted as a C^α^ node in the network. Notably, the experimental B-factor prediction was minimally affected by node placement at each atom, reinforcing the appropriateness of utilizing C^α^ residues. The spring constant, denoted as Gamma Value 1, was uniformly applied in constructing both models. The modes of GNM are obtained from the pseudo-inverse Kirchhoff contact matrix (Г− 1), expressed by the eigenvalues (λ_k_), in ascending order with the smallest, zero-valued eigenvalue omitted, and eigenvectors (u_k_). Notably, the summation excludes the initial eigenvalue (k = 1), which is equal to zero. (Haliloglu and Bahar, 1998)

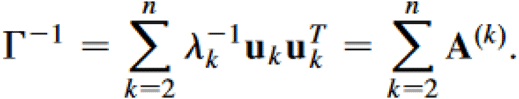

In this above equation, n is the number of residues of the protein, A^(k)^ is n × n matrix describing the contribution of the k^th^ vibrational mode to atomic fluctuations, superscript T indicates the transpose.

However, GNM lack directionality information, slow modes of motion of proteins are derived using the ANM method, allowed us to describe the directionality of the various modes. ANM modes are obtained from the Hessian matrix, expressed by harmonic potentials (Atilgan *et al*., 2001).

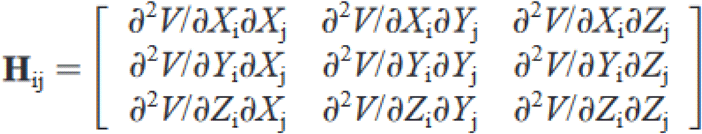

Here, V denotes harmonic potential and letters i and j represent residues, with X, Y, and Z denoting coordinates.

### Molecular Dynamics

We used the GROMACS package for MD simulation (Berendsen *et al.,* 1995, Hess et al., 2008). We constructed four setups to perform MD simulation: 1) open conformation with two Ca^2+^ ions; 2) open conformation without Ca^2+^ ion; 3) closed conformation with two Ca^2+^ ions; and 4) closed conformation without Ca^2+^ ion. We used the OPLS-AA/L all-atom force field (Kaminski *et al.,* 2001, Kulig et al, 2015) to generate the topology file. Then we defined the box type as cubic and set the distance between the protein and the edges of the box to 1.0 nm. We added 19 Na^+^ ions in open conformation with Ca^2+^, 23 Na^+^ ions in open conformation without Ca^2+^, 17 Na^+^ ions in closed conformation with Ca^2+^, and 21 Na^+^ ions in closed conformation without Ca^2+^ to neutralize the protein system. We ran energy minimization with the steepest descent algorithm for 50,000 steps to find the lowest potential energy conformation of the protein. The energy for the open conformation with Ca^2+^ ion was converged at -1.07607e^07^ kJ/mol at the 3634^th^ step, and that of the open conformation without Ca^2+^ ion was converged at -1.07673e^07^ kJ/mol at the 3942^nd^ step. That of the closed conformation with Ca^2+^ ion was converged at -4.18073e^06^ kJ/mol at the 2559^th^ step, and that of the closed conformation without Ca^2+^ ion was converged at -4.17851e^06^ kJ/mol at the 2555^th^ step. We performed NVT equilibration for 100 ps, followed by NPT equilibration for 100 ps to the system. Then, we ran all four setups in the production MD simulation for 200 ns. After the completion of the MD simulation, we processed the trajectory file to remove periodic boundary conditions, artificially induced rotations and translations, and centered the protein in the box. The processed trajectories were used for further analysis. We performed a solvent accessible surface area (SASA) analysis of four setups for localized structural changes (described below).

### Protein Domain Movement Analysis

We extracted the initial and final snapshots from the 200-ns trajectory PDB file using Python code. These two snapshots in PDB format were given on the Dynamics Domain (DynDom) server (http://dyndom.cmp.uea.ac.uk/dyndom/main.jsp) (Lee et al, 2003). The DynDom uses the k-means clustering algorithm to find the cluster of rotation vectors. These rotational vectors were used to calculate the domain motions of the protein. The domains in motion were colored red and the fixed domains were colored blue. The bending residues were colored in green, and the rotational vectors were represented as arrows.

### Principal Component Analysis

Principal Component Analysis (PCA) was used to obtain the correlated motions of the residues in the wild type (WT) open and closed conformations with Ca2+ bound and unbound. gmx_coavar, gmx_anaeig in GROMACS were used to perform PCA by constructing covariance matrix, eigenvectors and eigenvalues, which are used to calculate the correlated motions of the WT protein structures.

### Identification of solvent accessible residues

We have taken the absolute solvent accessibility value from the output of DSSP (Kabsch and Sander., 1983, Rost and Sander, 1994) and calculated the relative solvent accessibility. The formula for calculating relative solvent accessibility is

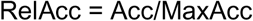

Where MaxAcc is maximal accessibility for the amino acids. Structures were run at 200 ns and the final atomic structures were used to generate the DSSP output file and calculate the solvent accessibility. We have used thresholds to distinguish two states initial and final state of our structure as stated in (Rost *et al*., 1994). Here, we considered a residue as Buried (B) if Relacc < 9%, and a residue as Intermediate (I) if Relacc = 9 – 36%, and a residue as Exposed (E) if Relacc >= 36%.

### Non-Synonymous SNPs of TG2 and their impact on protein stability

We collected the nsSNP data of the human TG2 gene from the SNPdbe database (Schaefer *et al.,* 2012) (https://www.rostlab.org/services/snpdbe/). There were 99 nsSNPs at the nucleotide sequence level and when the corresponding amino acid was checked there were redundant nsSNPs. Thus, we used 61 nsSNPs for both sequence-based and structure-based analyses to determine if the nsSNPs affect the stability of the protein.

We used sequence information-based prediction tools like PolyPhen-2 (Adzhubei *et al*., 2013), fathmm (Shihab *et al.,* 2012), PhD-SNP (Capriotti *et al.,* 2006), SuSPect (Yates et al., 2014), SNPs and GO (Calabrese *et al.,* 2009), I-Mutant 3.0 (Capriotti *et al*., 2005), SNAP (Bromberg *et al.,* 2007), and SIFT (Kumar *et al.,* 2009) to predict the structures of mutated TG2. These tools use sequences and SNPs as input to predict the effect or lack of effect of disease-associated mutations. The use of comparative physical and evolutionary conservation by PolyPhen2.0 (polymorphism phenotyping) (http://genetics.bwh.harvard.edu/pph2/) evaluates the potential impact of amino acid alterations on the structure and functionality of the protein (Adzhubei *et al*., 2013). A high-throughput web server called fathmm (http://fathmm.biocompute.org.uk/) uses hidden Markov models (HMMs) to determine the functional effects of coding changes (Shihab *et al., 2012*). PhD-SNP (https://snps.biofold.org/phd-snp/phd-snp.html) is a support vector machine (SVM)-based model that distinguishes between mutations linked with disease and is based on the profile of sequence conservation (Capriotti *et al.,* 2006). SuSpect (http://www.sbg.bio.ic.ac.uk/suspect/) predicts the phenotypic implications of missense variants using sequence, structure, and systems biology methods (Yates et al., 2014). The SNPs&GO (https://snps-and-go.biocomp.unibo.it/snps-and-go/) tool uses an SVM classifier to identify disease-related variants using a protein sequence profile and the functional data expressed in gene ontology terms (Calabrese *et al.,* 2009). The SNAP (screening for non-acceptable polymorphism) tool (https://www.rostlab.org/services/SNAP/) is based on neural networks (Bromberg *et al.,* 2007). The SIFT tool uses a homology sequence to determine if the mutation affects the function of a protein or not by assigning a score for damage prediction (Kumar *et al.,* 2009). The I-Mutant 3.0 (http://gpcr2.biocomp.unibo.it/cgi/predictors/I-Mutant3.0/I-Mutant3.0.cgi) tool is based on the SVM method to predict damage from single point mutations (Capriotti *et al*., 2005).

We used the following structure-based protein stability procedure tools: ERIS (Yin *et al.,* 2007), INPS (Fariselli et al., 2015), Dynamut (Rodrigues *et al.,* 2018), Maestro (Laimer *et al*., 2015), DUET (Pires et al., 2014), and I-Mutant 3.0 (Capriotti *et al*., 2005). All the structure-based calculations calculate the free energy by estimating Gibbs free energy. The Gibbs free energy is calculated to predict the stability of proteins. The ΔΔG is calculated by estimating the difference between the ΔG of mutant structure and the ΔG of wild-type structure (Gilis and Rooman, 1997). The unit of ΔΔG is kcal/mol.

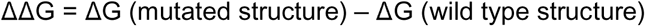

The positive ΔΔG value indicates that the structure is unstable, and the negative value indicates that the structure is stable. The ERIS server (https://dokhlab.med.psu.edu/eris/login.php) is an automated tool that predicts the protein stability induced by mutation. The DynaMut (http://biosig.unimelb.edu.au/dynamut/) is a web server that integrates graph-based signatures and normal mode dynamics to generate a consensus prediction of the impact of a mutation on protein stability (Rodrigues et al., 2018). The INPS (Impact of Non-synonymous mutations on Protein Stability) (https://inpsmd.biocomp.unibo.it/inpsSuite/default/index) algorithm, which is based on the SVM method, is trained to predict the change in Gibbs free energy resulting from single-point mutations in protein sequences. Maestro (https://pbwww.services.came.sbg.ac.at/maestro/web) is a web-based tool that uses statistical scoring functions and machine learning methods like neural networks, SVM, and multiple linear regression (Laimer et al., 2015). DUET (http://structure.bioc.cam.ac.uk/duet) uses the statistical potential energy function method to predict the free energy of the proteins (Pires et al., 2014). The I-Mutant 3.0 (http://gpcr2.biocomp.unibo.it/cgi/predictors/I-Mutant3.0/I-Mutant3.0.cgi) tool is based on the SVM method to predict stability for single point mutations (Capriotti et al., 2005).

## Results and Discussion

There were nine crystal structures available in the Protein Data Bank (PDB) (wwPDB consortium, 2019) with the query "*tgm2*" and source organism “*Homo sapiens*.” Among these we found five structures in the closed conformation and four structures in the open conformation. Both the conformations are bound with either (GTP/GDP/ATP) or a peptide. The crystal structures that were analyzed in this study are listed in Supplementary Table 1 and Table 2.

99 non-synonymous SNPs (nsSNPs) were reported for human TG2 in SNPdbe (Schaefer *et al*., 2011) obtained by computational or experimental methods. After reducing redundancies, the number of nsSNPS was reduced to 61. We then looked at their spatial location and proximity to active site area and surface accessibility.

### Coarse-grained modeling of TG2

For building coarse grained models for each structure, we calculated total number of residues and total number of modes from slowest to fastest using ProDy (Zhang *et al.,* 2021). To validate the GNM models as significant and can be used for further analysis, the mean square fluctuation (MSF) values for all structures were calculated and correlated with the experimental B-factors (Table 1). The B-factor or temperature factor or Debye-Waller is a measure of the flexibility of protein structures. The smaller B-factor values indicate the higher stability of a residue/atom, and the higher the value indicates the more flexibility. It has been reported that the correlation between B-factor and MSF should be greater than 0.58 for an ENM model to be considered significant (Kundu *et al*., 2002). As shown in Table 1, six out of nine PDB structures had a MSF vs. B-factor correlation coefficient greater than 0.58. The plots of correlation are shown in Figure 2. A higher correlation of B-factor vs MSF was observed in 3S3S with the value of 0.78 (Figure 2C) in the open conformation structures. Similarly in the closed conformation structures we observed 4PYG with the higher correlation of B-factor vs MSF with the value of 0.69 (Figure 2F).

**Figure 2:**
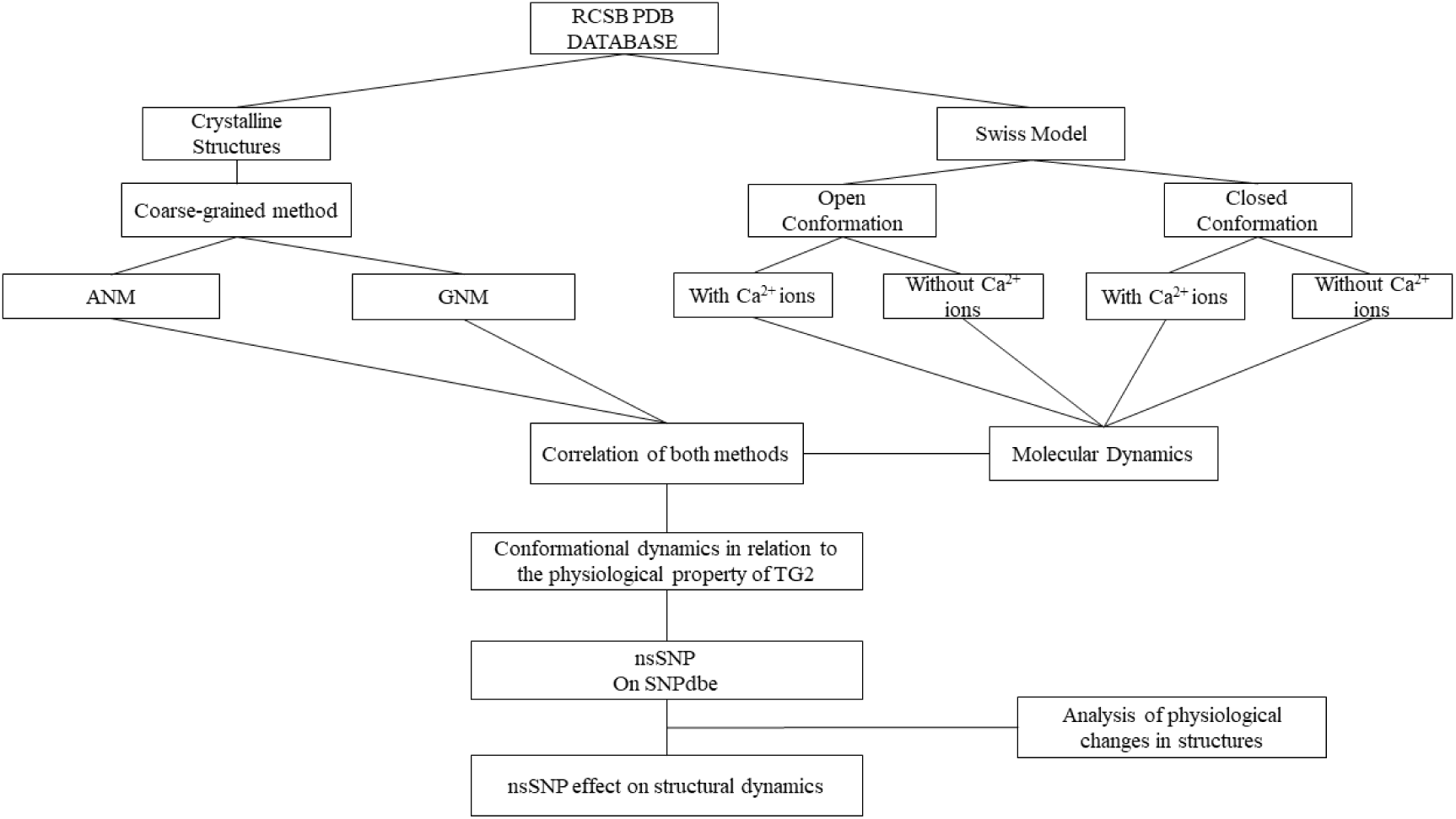
**The workflow adopted for this study.**

**Table 1:**
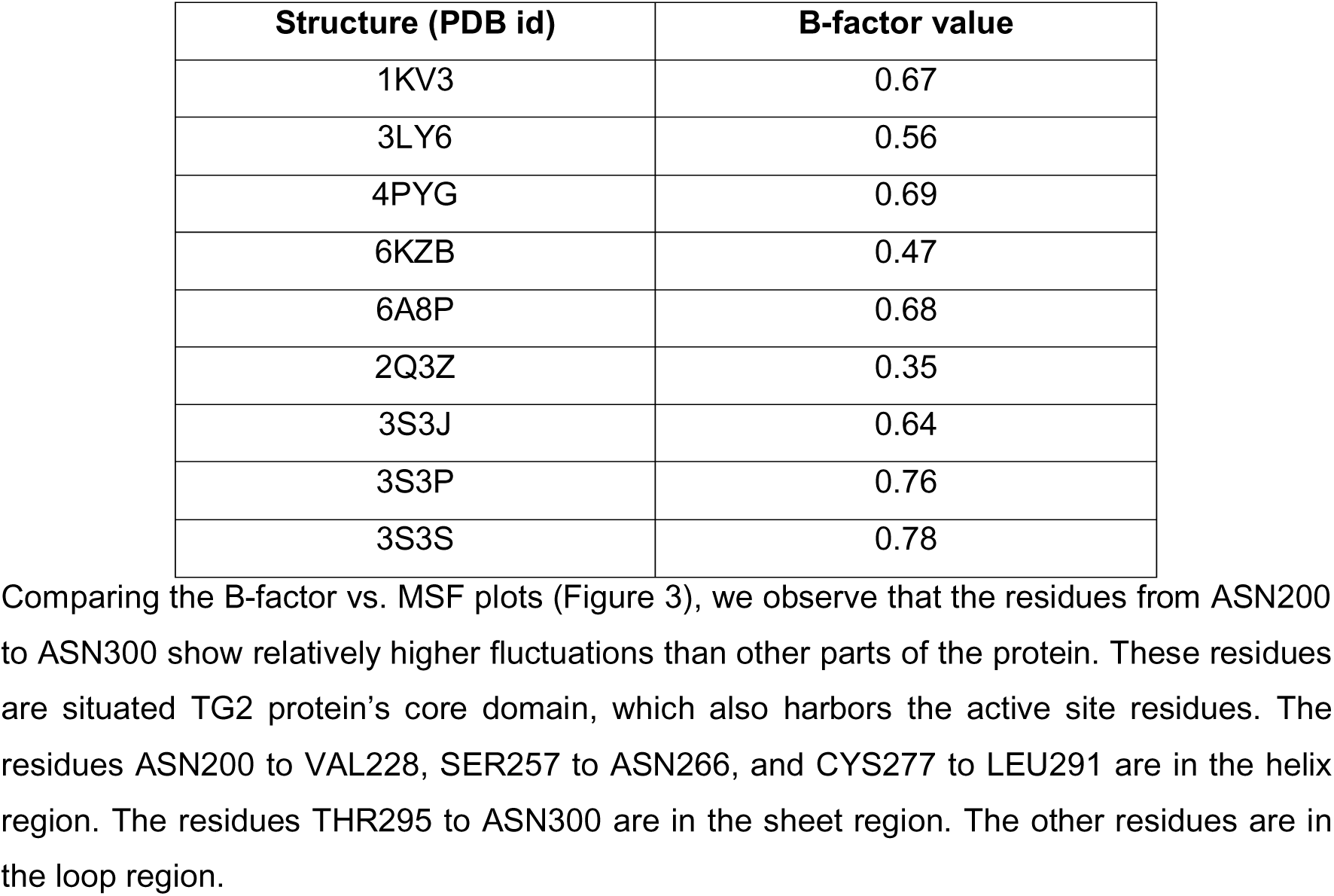
MSF versus B-factor correlation with corresponding PDB structures.

| Structure (PDB id) | B-factor value |
| --- | --- |
| 1KV3 | 0.67 |
| 3LY6 | 0.56 |
| 4PYG | 0.69 |
| 6KZB | 0.47 |
| 6A8P | 0.68 |
| 2Q3Z | 0.35 |
| 3S3J | 0.64 |
| 3S3P | 0.76 |
| 3S3S | 0.78 |

### Intra-Molecular Dynamics

GNM cross-correlation maps for the first 10 slowest modes were plotted individually to identify the regions that are either positively or negatively correlated in the protein’s dynamics (Figure 4). Here, we compared the first three slowest modes of both open and closed conformations. The cross-correlation map is significantly correlated with MSF. The red color block indicates the positively correlated and blue color block indicates the negatively correlated regions within the protein. The cross-correlation values range from −1.0 to +1.0.

**Figure 3:**
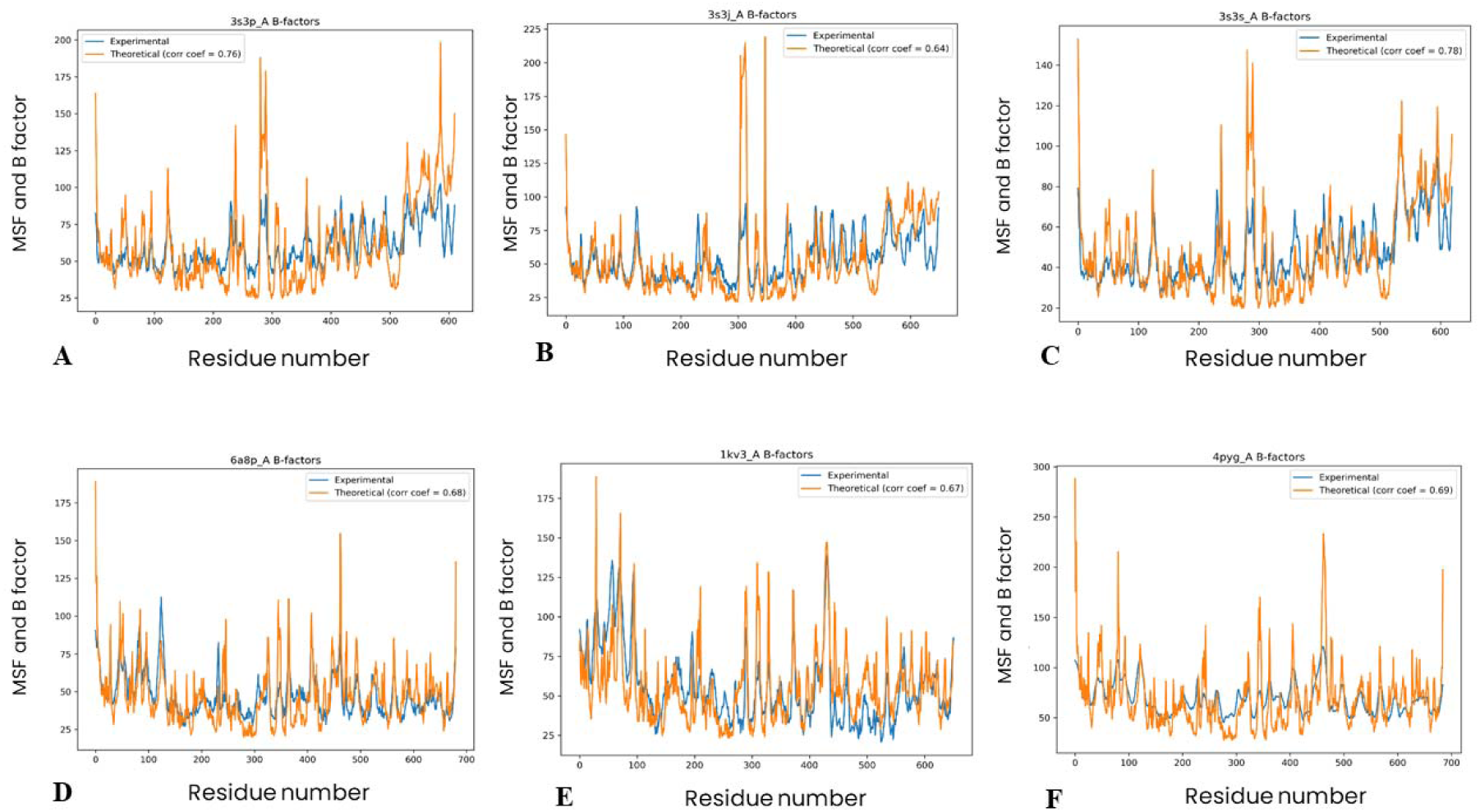
Mean Square Fluctuation (MSF) versus B-factor correlation. A, B, and C are the correlation plots for the open conformation (PDB id: 3S3P, 3S3J, and 3S3S, respectively). D, E, and F are the correlation plots for the closed conformations (PDB id: 6A8P, 1KV3, and 4PYG, respectively). The orange line indicates the MSF and the blue line indicates experimental B-factor.

**Figure 4:**
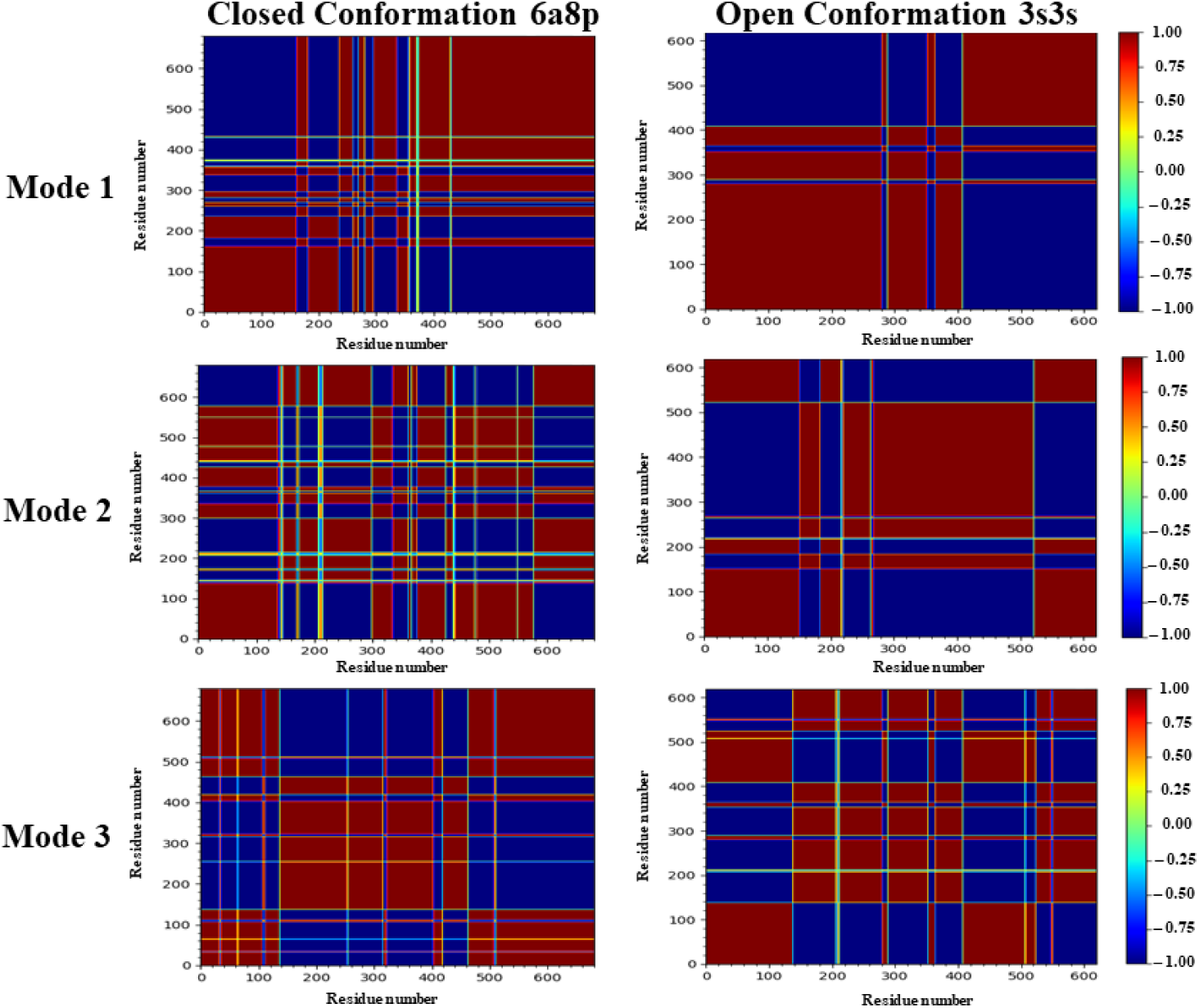
First three slowest mode of cross correlation plots from GNM. The red color block indicates the positively correlated and the blue color block indicates the negatively correlated regions.

In open conformation, first three slowest modes of 6A8P, 1KV3, 4PYG have the same interdomain flexibility. Similarly, to the closed conformation 3S3S, 3S3J, 3S3P have the same interdomain flexibility for the first three slowest modes (Supplementary Figure 1 and Supplementary Figure 2). So, we considered the crystal structure with the highest MSF vs. B-factor correlation value in both conformations. In the open conformation, 3S3S has the highest correlated B-factor with the value of 0.78. Similarly, in the closed conformation 4PYG has the highest correlated B-factor value of 0.69. We compared the cross-correlation maps of these two protein structures. For the 1^st^ slowest mode, the intramolecular core domain is positively correlated and have significance changes in domain dynamics for both open and closed conformations (Figure 3 Mode 1). Two positively correlated blocks show inter domain flexibility in mode 1. Compared to the open conformation of mode 1, that of the closed conformation is highly fragmented and shows low correlation. The lower correlation in the open conformation informs about the low level of interaction.

For the second slowest mode, cross-correlation maps are more delineated than the first mode of the closed conformation. The closed conformation has more fragmented dynamic regions than those of the open conformation (Figure 3 Mode 2). Four blocks show interdomain flexibility in the closed conformation. There are three blocks and have interdomain flexibility in the closed conformation.

For the third slowest mode, cross-correlation maps are more delineated than the second mode of the open conformation. The open conformation has more fragmented dynamic regions than closed conformation (Figure 3, Mode 3). Three blocks show interdomain flexibility in the closed conformation. There are four blocks and have interdomain flexibility in the closed conformation. These regions of correlation enable the protein to contain intrinsic fluctuations, which most likely would enable interaction with ligands and substrates.

### Directionality in the dynamics

The first three slowest modes derived from ANM revealed that some regions were more relatively flexible and dynamic than the other parts of the structure (Figure 5). The N-terminal β-sandwich domain has relatively more flexibility compared to all other domains. These dynamic patterns were observed in both conformations. The dynamics around the active site residues were more subdued, implying that these fluctuations do not affect the ligand interactions.

**Figure 5:**
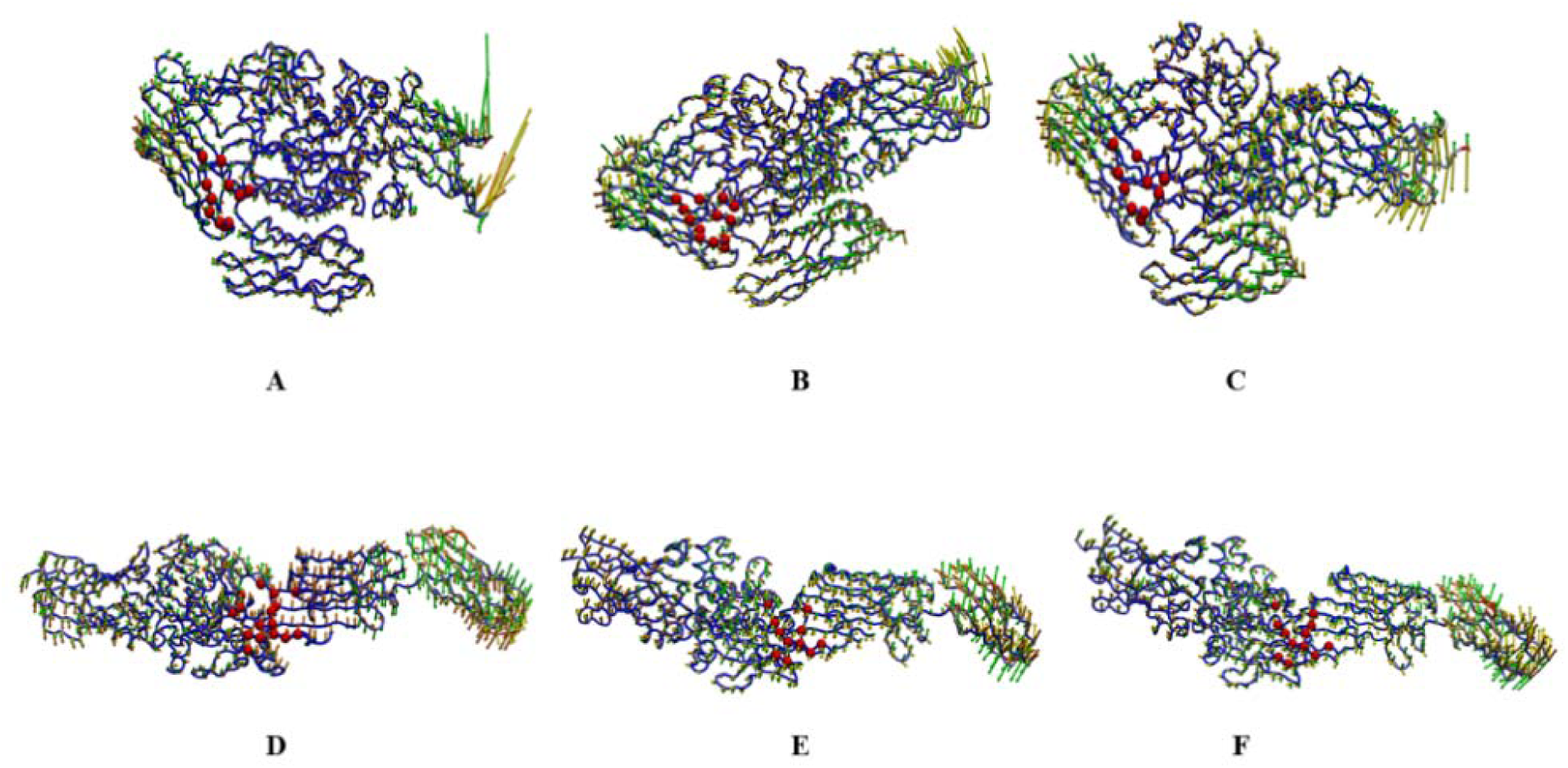
First three slowest mode plots from ANM. The highly dynamic regions are showed in red and least dynamic region are showed in blue. The orange, yellow, and green arrows represent first, second, and third modes, respectively. The red sphere indicates the active site residues. A, B, and C are 1KV3, 4PYG, and 6A8P, respectively. D, E, and F are 3S3J, 3S3P, and 3S3S, respectively.

### Principal Component Analysis

PCA analyses was done for WT closed with Ca2+ ions, WT closed without Ca2+ ions, WT open with Ca2+ ions, WT open without Ca2+ ions (Figure 6). The plots were generated using backbone atoms and plotted the PC1 against PC2. Interestingly, it is depicted from the plots that the wild type (WT) closed conformations has higher number of distinct cluster patters.

**Figure 6:**
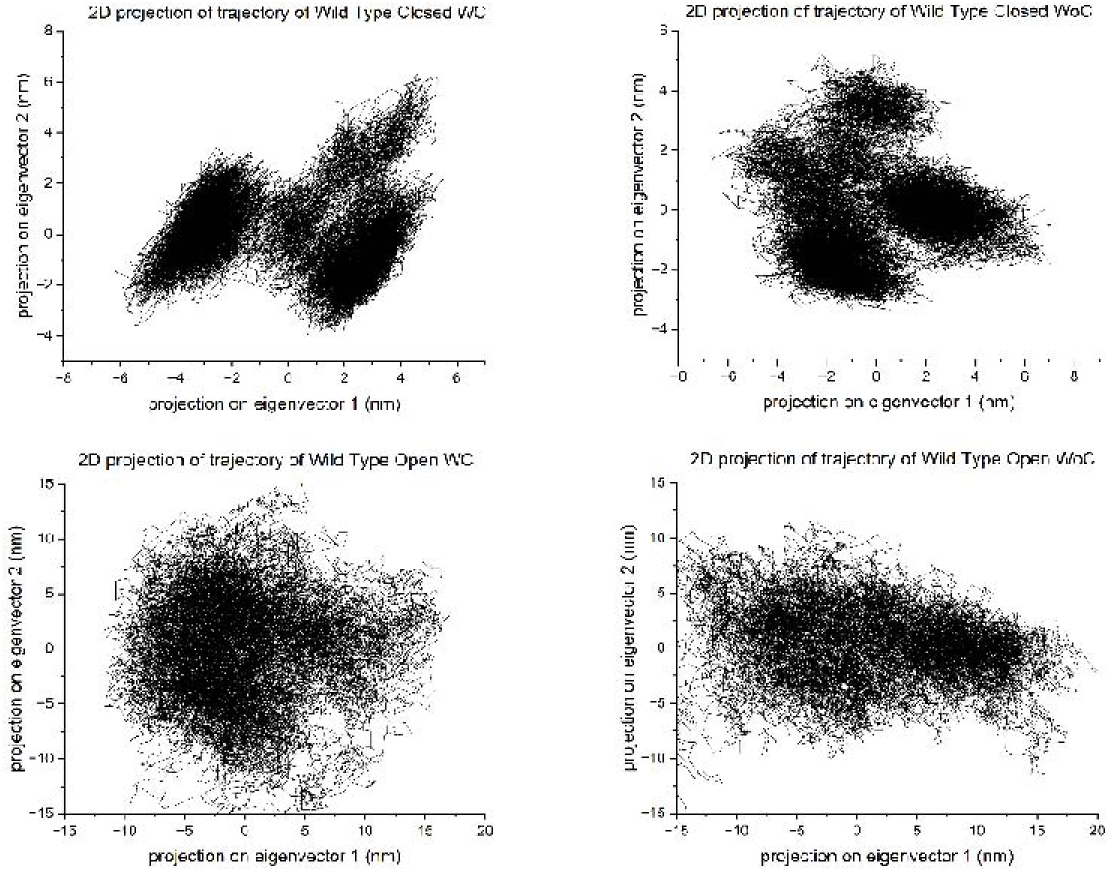
Two dimensional plots from the results of PCA for WT open and closed conformations.

### Global cross correlation

The global cross correlation modes were obtained from ANM (Figure 7), where negative correlation are colored blue and positive correlation are colored red. Here, we observe that there are relatively more negatively correlated regions than those that are positively correlated. The “all modes” of the open and closed conformations have similar behaviors (Figure 5). In the open conformation, there are only three regions that show flexibility. This imply that closed conformation is rigid during dynamics. Based on the cross-correlation amplitudes between the two conformations (Figure 5), the positive correlation in open conformation region may point to TG2’s interaction region with peptides (i.e., gluten).

**Figure 7:**
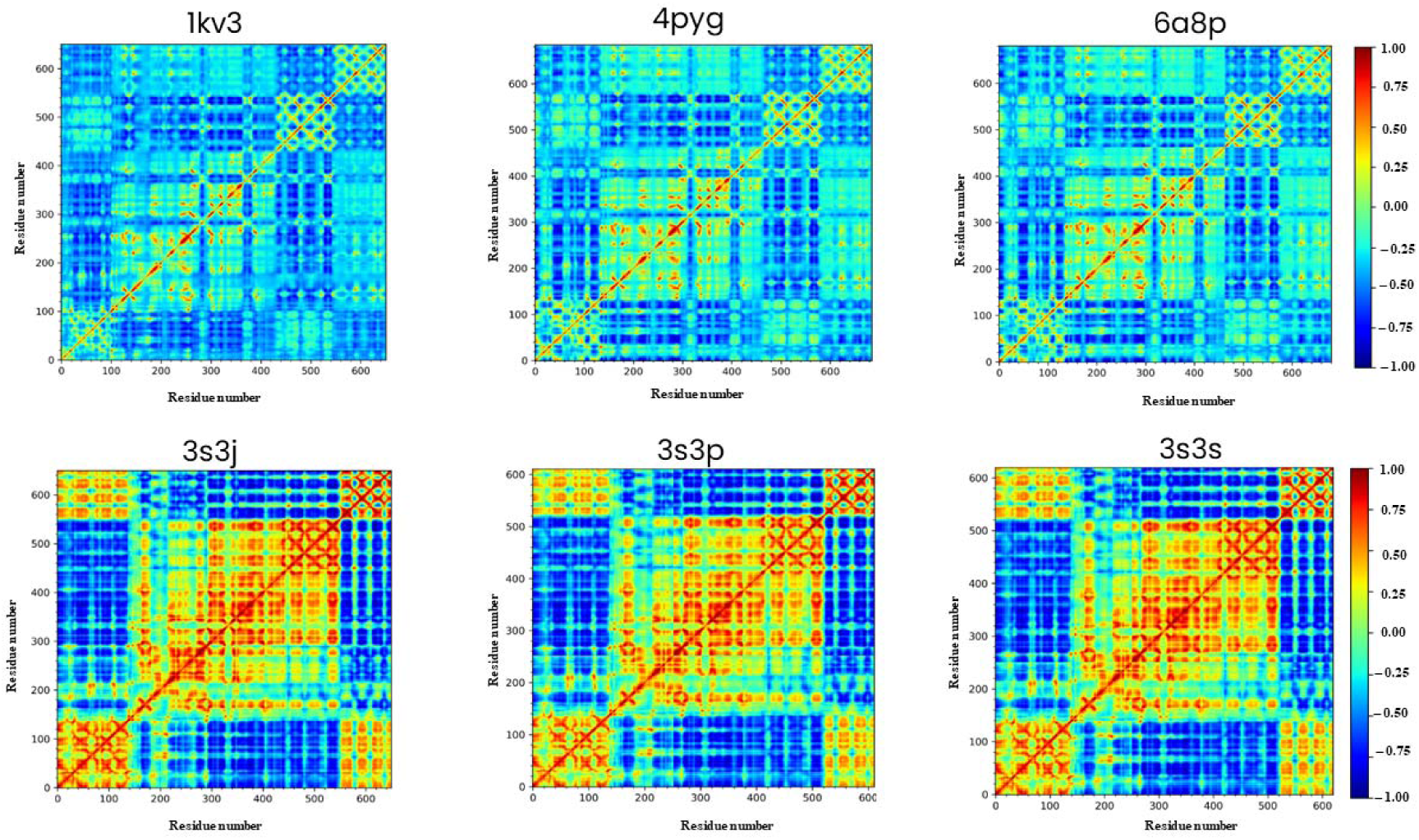
Global cross correlation map obtained from ANM, where the top row shows closed conformations (1KV3, 4PYG, and 6A8P) and the bottom row open conformations (3S3J, 3S3P, and 3S3S). The blue color indicates negative correlation. The red color indicates positive correlation.

**Figure 8:**
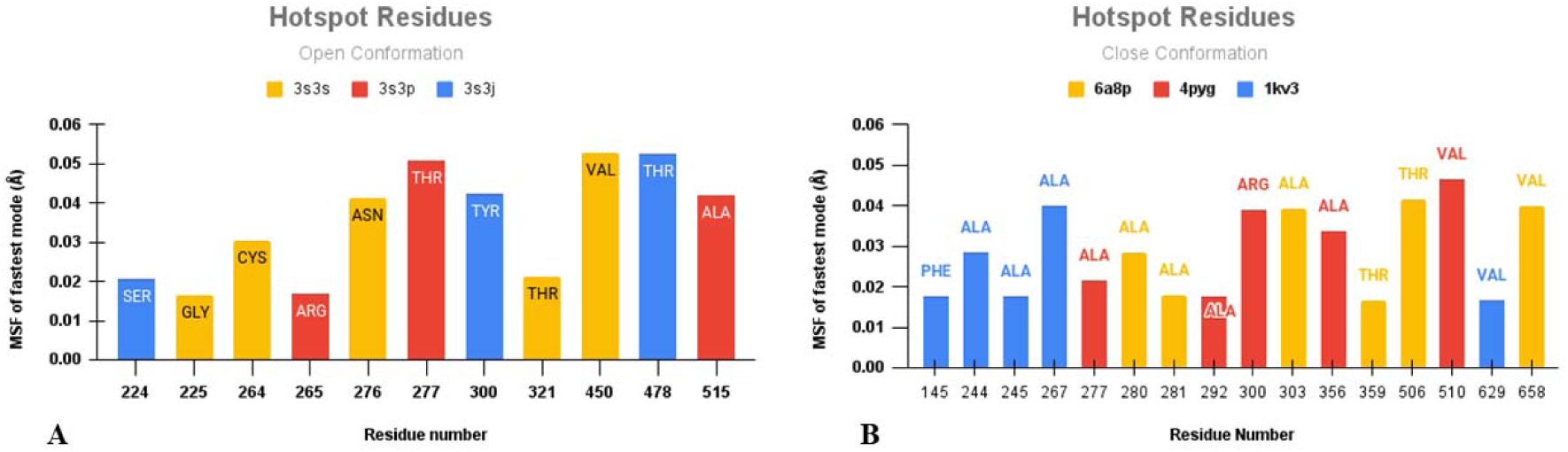
Mean Square Fluctuations of hotspot residues for the five fastest modes. Open conformation [A] and close conformation [B]. Color indicates the results for different structures.

### Hotspot regions

Hotspot residues are specific amino acid residues in the protein-protein interaction interface that make a significant contribution to the binding affinity and stability of the protein complex (Keskin et al., 2005, Halperin et al., 2004, Ma et al, 2003, Demirel et al., 1998, Haliloglu et al., 2005). These residues are often crucial for the formation of specific protein-protein interactions and play a central role in the recognition and binding between proteins. The residues in hotspot region may participating in ligand interaction. we hypothesize that these hotspot residues have a role in TG2’s function. Further, we analyzed the five fastest modes (Figure 6) for both the conformations of TG2. The fastest modes show biological significance in the binding region. The highly fluctuating residues at fast modes play important role in interaction, stability, and protein functions (Demirel et al., 1998).

The hotspot residues in the open conformation are near the active site region of TG2 (Figure 9). In the closed conformation, the hotspot regions are distant from the active site residues (Figure 10). In 1KV3, the distance between the hotspot region and active site region is 8.0 Å. In 4PYG, the distance between the hotspot region and active site region is about 8.8 Å. In 6A8P, the distance between the hotspot region and active site region is about 8.2 Å. This relative closeness of the hotspot region to the active implies that the hotspot residues in open conformation is highly likely in the peptide binding.

**Figure 9:**
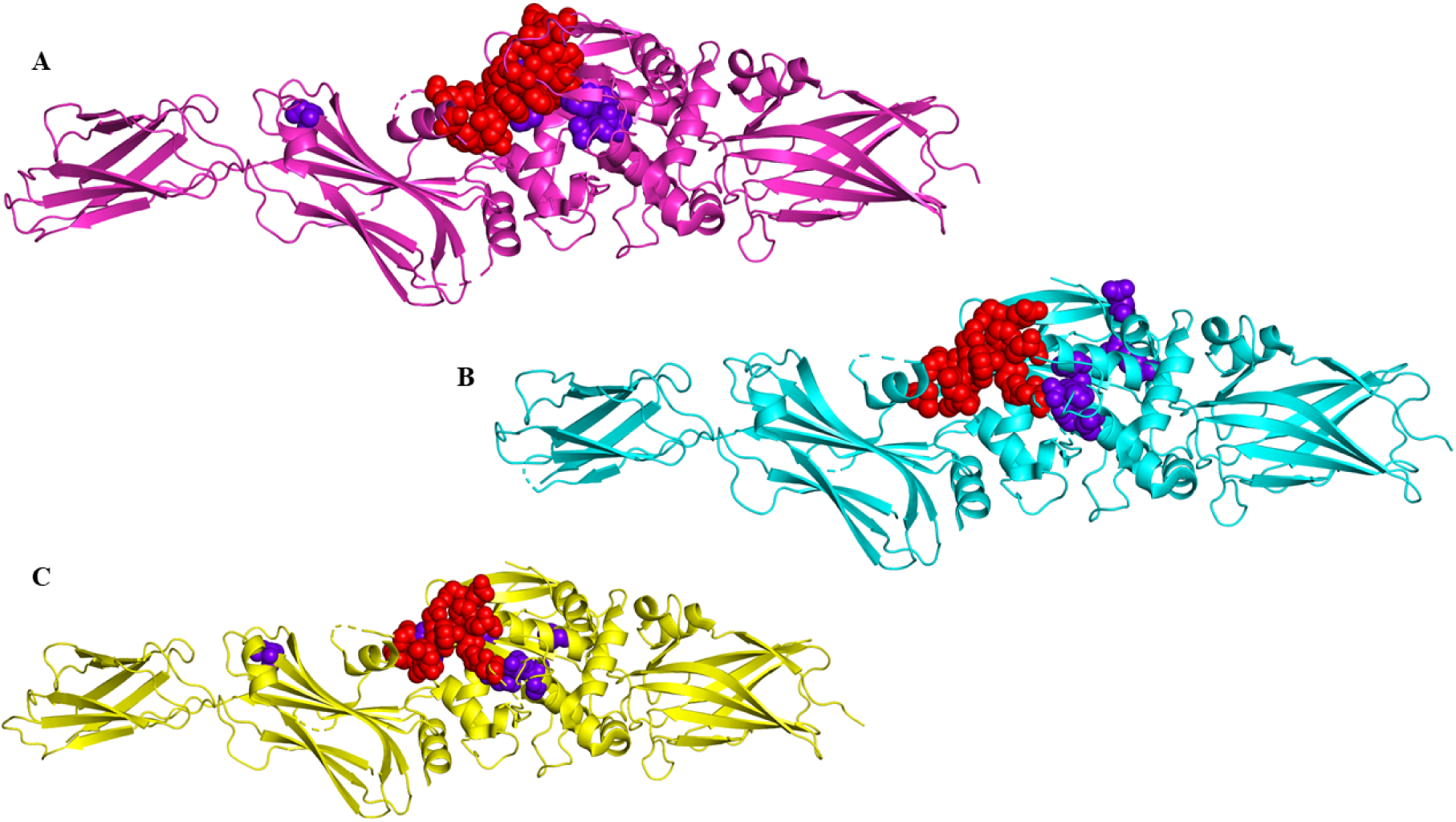
Spatial Proximity of hotspot residues with active site residues (open conformation). A, B, and C are 3S3J, 3S3P, and 3S3S. Red sphere indicates active site residue and purple indicates hotspot residues.

**Figure 10:**
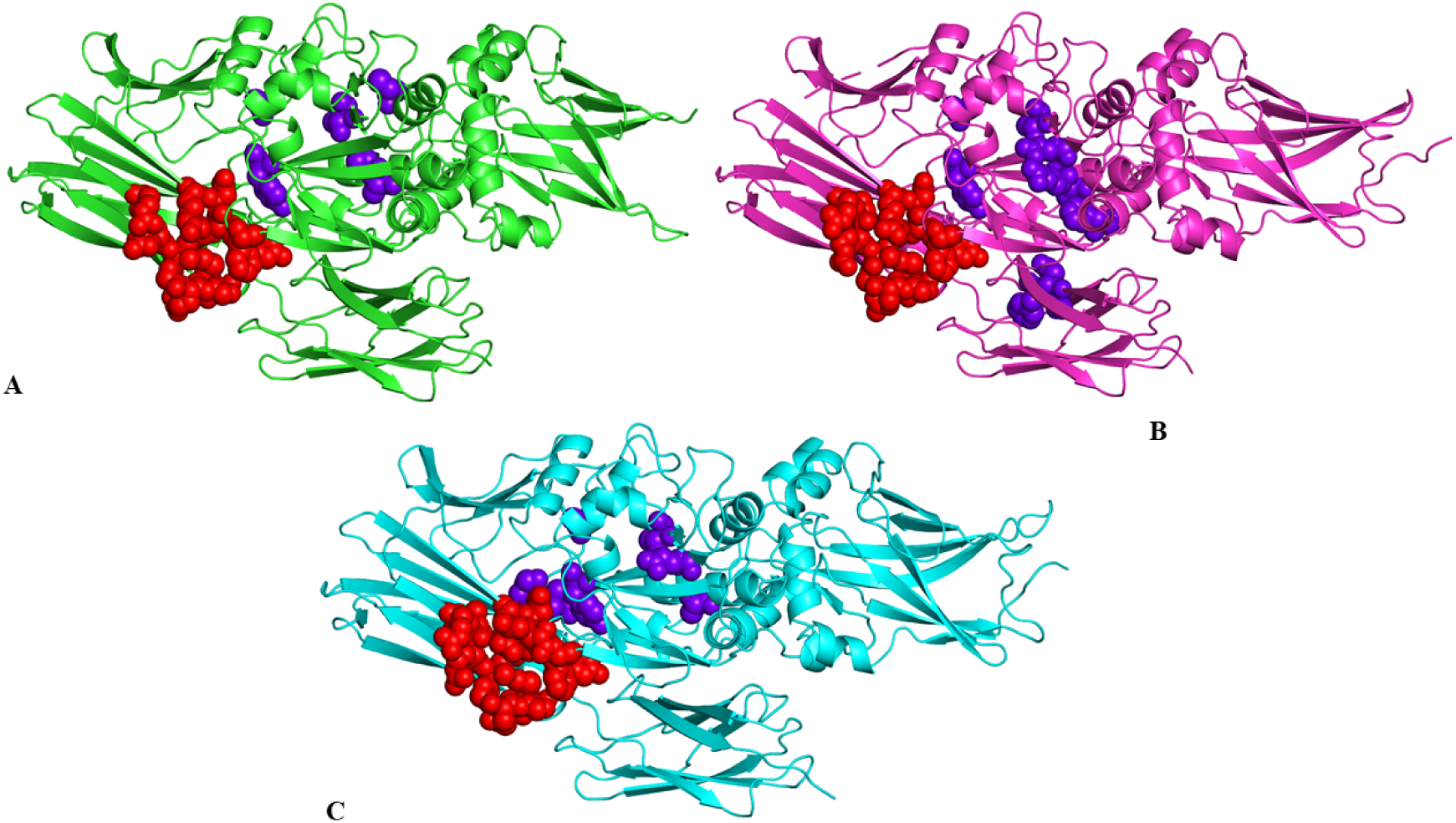
Spatial Proximity of hotspot residues with active site residues (close conformation). A, B, and C are 1KV3, 6A8P, and 4PYG. Red sphere indicates active site residue and purple indicates hotspot residues.

### Molecular dynamics simulation of TG2

For MD simulations, we used homology modelled proteins of TG2 in both conformations, as described in Methods section (Supplementary Figure 3). We performed 200 ns of MD simulation for the four setups and the results indicate significant conformational changes with and without calcium.

The initial and final snapshot of the 200 ns trajectory of open conformation (Figure 11) and closed conformation (Figure 11) were compared and the root mean square deviations (RMSD) were calculated between two structures. For the open conformation with a Ca^2+^ ion and without a Ca^2+^ ion, the RMSD differences were 2.342 Å and 2.511 Å (Figure 9A), respectively. In the open conformation without a Ca^2+^ ion, the β2 domain moved 37.9 Å from the initial structure during the simulation (Figure 9B).

**Figure 11:**
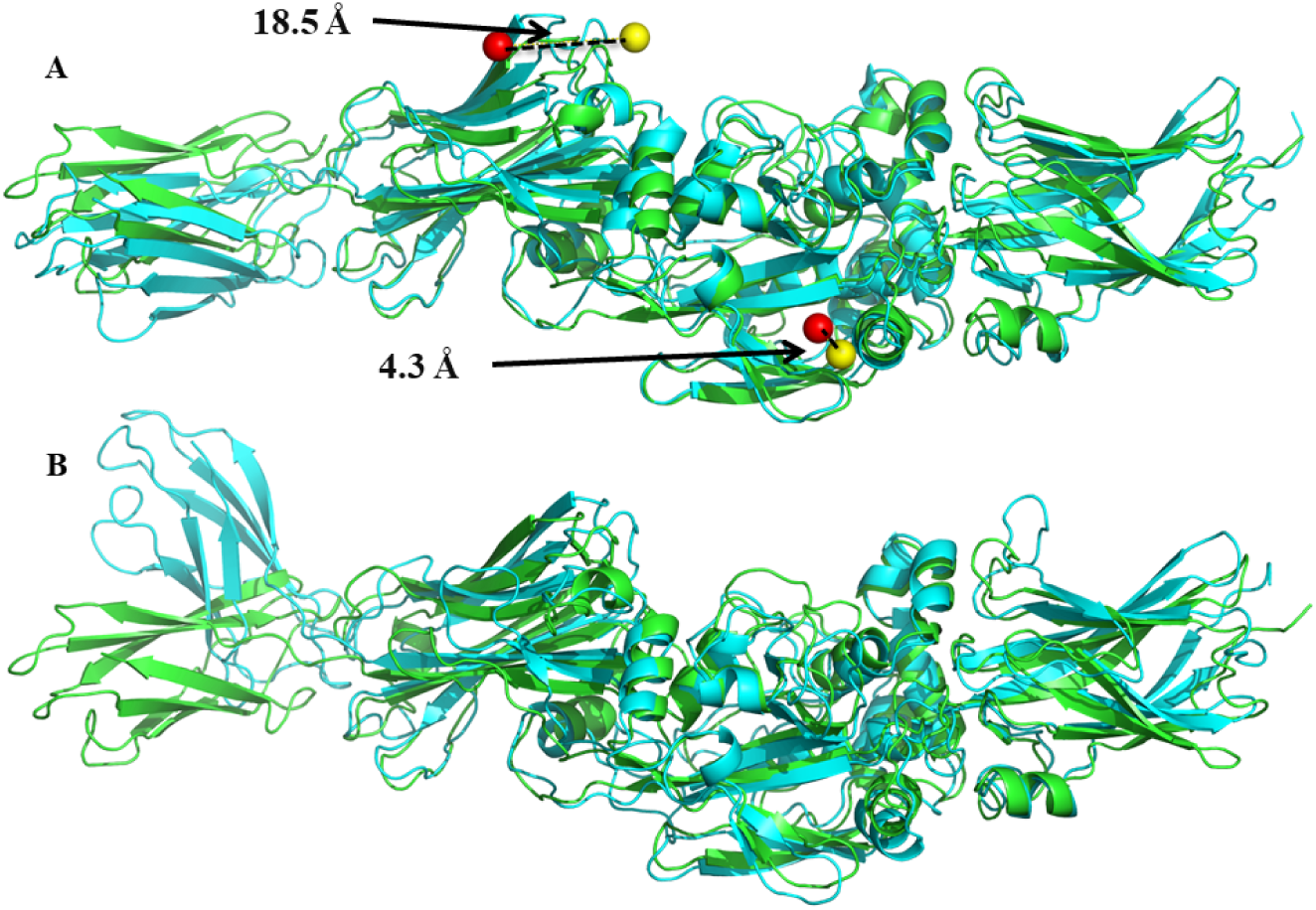
Superposed Structures of the open conformation of initial and final 200^th^ ns structure. The structure colored in green is the initial conformation (0 ns) with the corresponding Ca^2+^ ions shown as red spheres. The structure colored in cyan is the final conformation (200ns) with the corresponding Ca^2+^ ions shown as yellow spheres.

For the closed conformation with Ca^2+^ ions and without Ca^2+^ ions, the RMSD differences were 1.359 Å and 1.211 Å (Figure 12A) respectively. The two Ca^2+^ ions in the open conformation of final structure moved 18.5 Å and 4.3 Å from their initial positions. The two Ca^2+^ ions in the closed conformation of final structure moved 6.7 Å and 5.8 Å from their initial positions. In the closed conformation without Ca^2+^ ions, there were no significant domain movements compared to the structure at 0 ns (Figure 12B).

**Figure 12:**
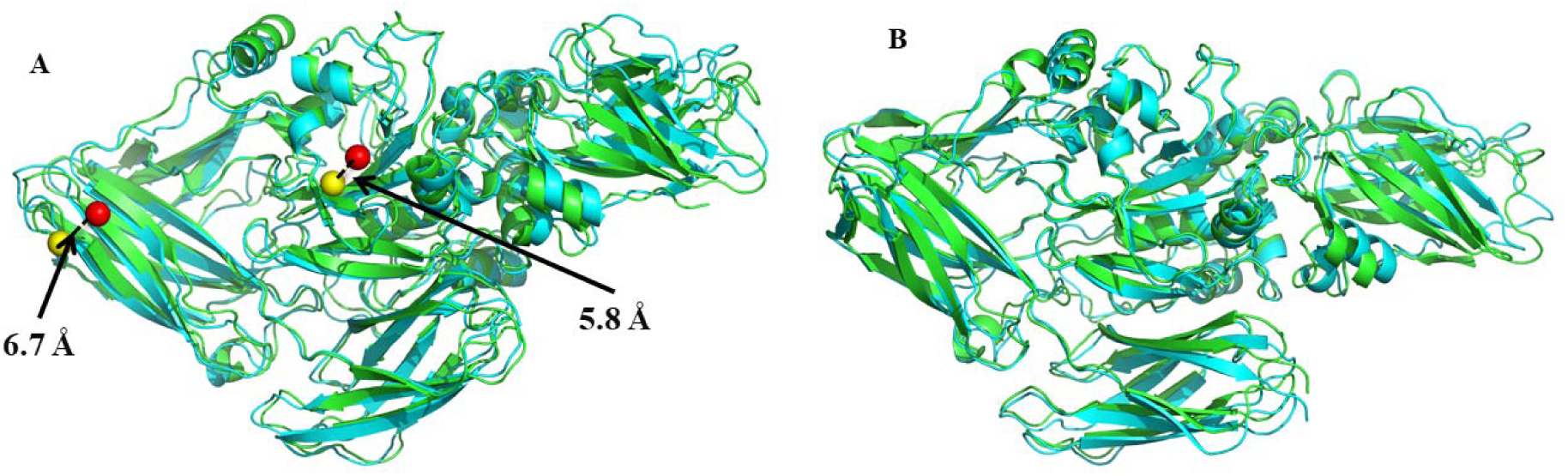
Superposed structures of the closed conformation of initial and final 200^th^ ns structure. The structure colored in green is the initial conformation (0 ns) with the corresponding Ca^2+^ ions shown as red spheres. The structure colored in cyan is the final conformation (200ns) with the corresponding Ca^2+^ ions shown as yellow spheres.

### Domain movement analysis in the four setups

We used DynDom (Lee *et al.,* 2003) for analyzing the domain movement based on MD trajectories. In the open conformation with Ca^2+^ (Figure 13A), the β2 domain moved in an upward direction, whereas in the open conformation without Ca^2+^ (Figure 13B) the β2 domain moved in a downward direction as represented by the rotational vectors. In all the four setups, the β2 domain showed the most mobile domain movement as compared to other domains (Figure 13). In closed conformations with Ca^2+^ and without Ca^2+^, the β2 domain was observed moving in a downward direction, as represented by the rotational vectors (Figure 13C and 13D). The rotational angles of four setups are shown in Table 2, where a higher rotational angle value means a larger conformational change in the structure. Thus, the open conformation without Ca^2+^ shows larger conformational change among four setups.

**Figure 13:**
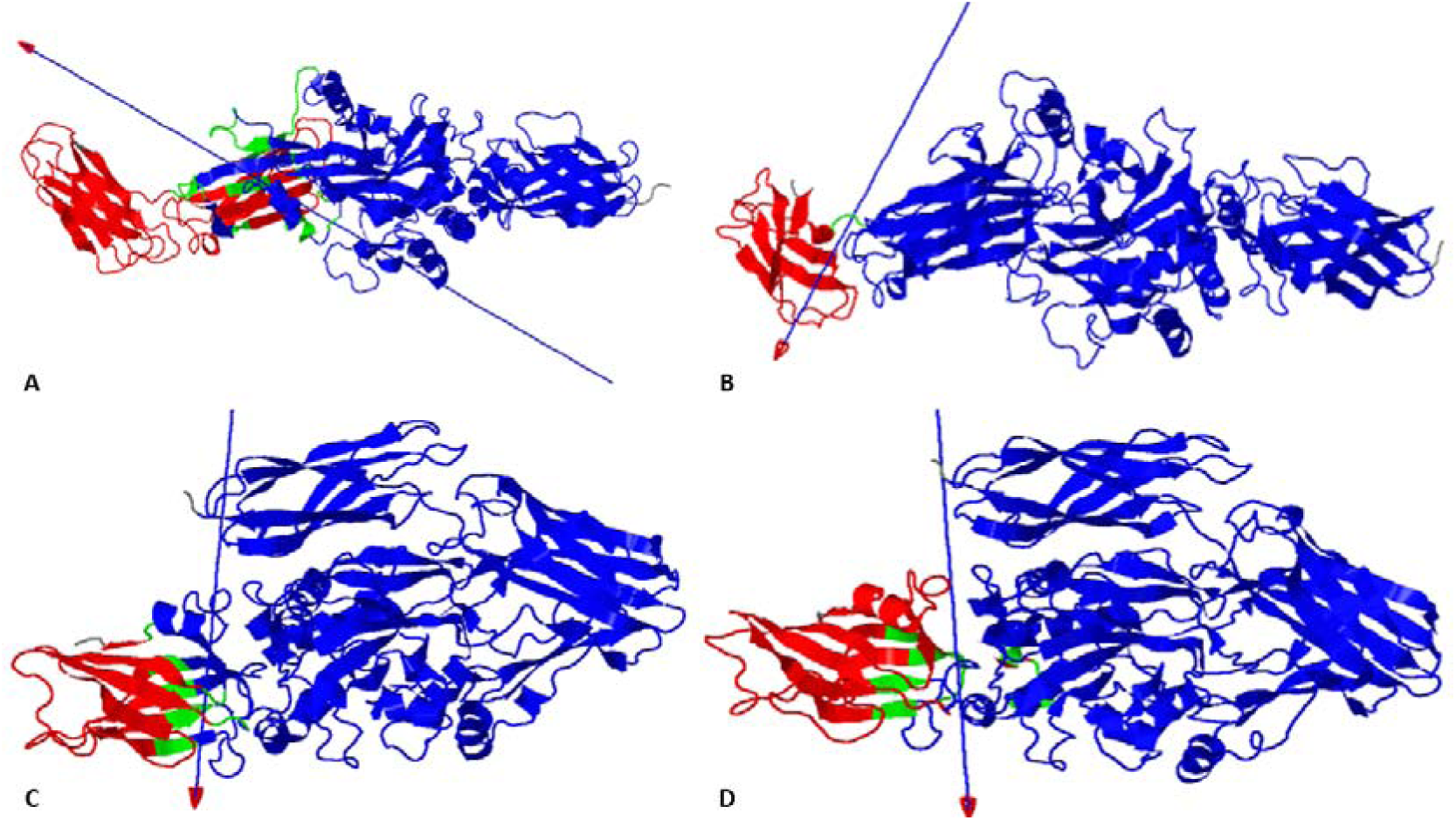
Domain motion in the four MD setups. A) domain motion of the open conformation with Ca^2+^ between initial and final 200^th^ ns structure, B) domain motion of the open conformation without Ca^2+^ between initial and final 200^th^ ns structure, C) domain motion of the closed conformation with Ca^2+^ between initial and final 200^th^ ns structure and D) domain motion of the closed conformation without Ca^2+^ between initial and final 200^th^ ns structure. Green, red, and blue represent bending residues, moving domain, and fixed domain, respectively. The arrow represents rotational angle vector.

**Table 2:** Rotational angle of derived from the first and final coordinated of the four MD setups.

| Setups | Rotational angle<br>(deg) |
| --- | --- |
| Open conformation with $\text{Ca}^{2+}$ | 16.4 |
| Open conformation without $\text{Ca}^{2+}$ | **66.1** |
| Closed conformation with $\text{Ca}^{2+}$ | 21.4 |
| Closed conformation without $\text{Ca}^{2+}$ | 18.0 |

### Solvent Accessible Surface Area (SASA) of active site residues in the open and closed conformations

The active sites of the protein were derived from PDBsum (Laskowski, 2022) to check if local changes took place in the trajectory. The Solvent Accessible Surface Area (SASA) in the open conformation was higher than that of the closed conformation (Figure 14). From 30 ns to 80 ns, a higher SASA is observed in the open conformation with Ca^2+^ compared to the open conformation without Ca^2+^. Closed conformations with Ca^2+^or without Ca^2+^had no significance difference in SASA.

**Figure 14:**
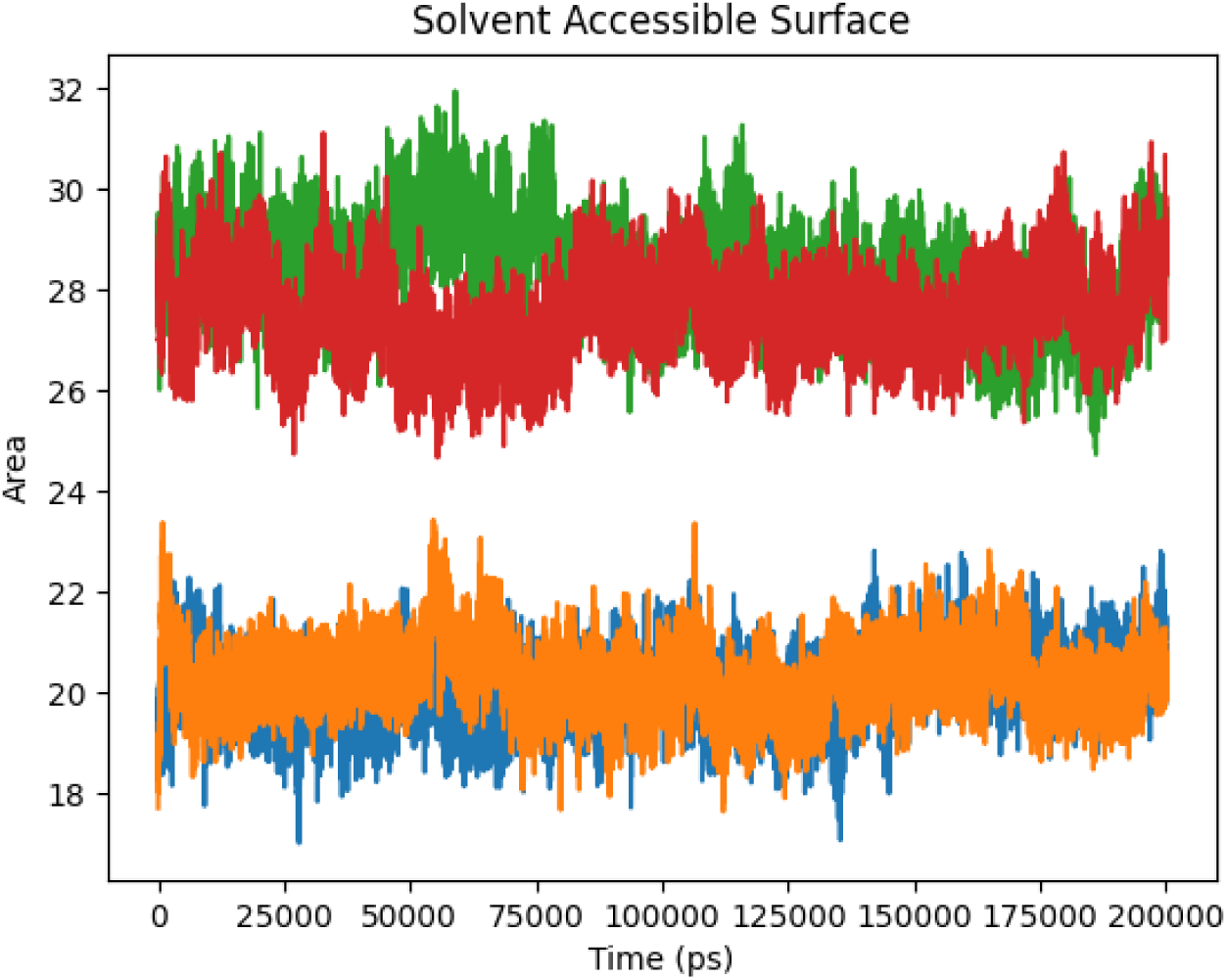
Solvent Accessible Surface Area (SASA) plot. Blue, orange, green, and red represent closed conformation with Ca^2+^ ion, closed conformation without Ca^2+^ ion, open conformation with Ca^2+^ ion and open conformation without Ca^2+^ ion.

**Figure 15:**
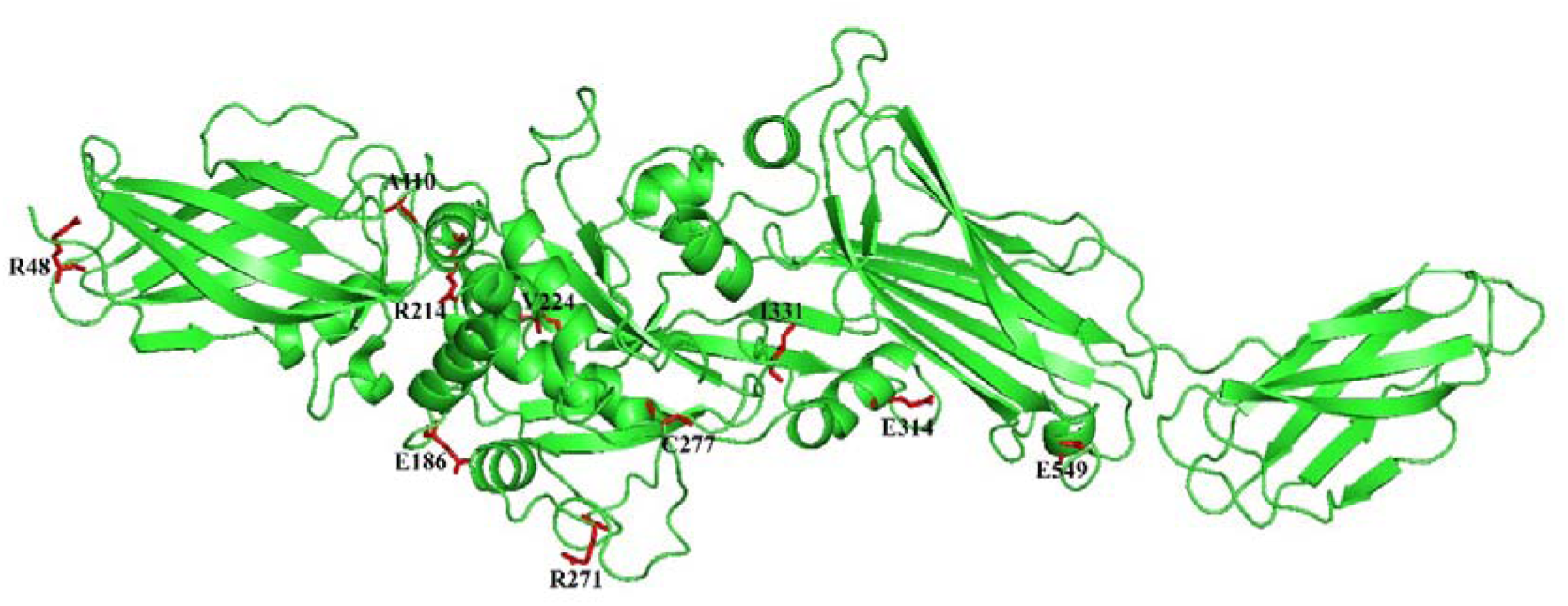
**Top 10 SNPs (red colored stick representations) positions marked in the three-dimensional TG2 structure in open conformation**

### Non-synonymous single nucleotide polymorphisms (nsSNPs) Impact Prediction Analysis

Amino acid substitutions, or non-synonymous single nucleotide polymorphisms (nsSNPs), can precipitate modifications in protein-protein interactions, stability, and enzymatic activity (Singh et al, 2020). The impact of nsSNPs on protein structure is highly variable, contingent on the specific amino acid alteration and its location within the protein. Such changes can provoke shifts in charge, polarity, size, hydrophobicity, or other physicochemical attributes of amino acids, which can lead to structural adjustments (Ng and Henikoff, 2001). Large inter-domain distances or amino acid substitutions can influence protein stability, folding, interactions, and overall structure-function correlations. Active site residues are vital for enzyme catalytic function, and proximal nsSNPs can interfere with interactions between active site residues and substrate molecules, which can lead to altered catalytic activity, substrate binding affinity, or specificity (Thusberg and Vihinen, 2009). For example, calcium ions have a fundamental role in protein structure and function, and nsSNPs near a calcium ion binding site can disrupt the coordination of calcium ions, potentially affecting protein stability and folding. The distance between domains and the presence of nsSNPs can potentially influence the structure of proteins. If the domain separation during dynamics is too large, it can lead to increased flexibility and potential impacts on the protein’s overall structure and function. On the other hand, if the domain separation is too small, steric conflicts between domains can arise, leading to structural restrictions and potential loss of protein functionality.

In this study, we used multiple prediction tools for consensus ranking approach for classifying the nsSNPs with the most substantial effects on sequence and destabilizing impacts on structure. For sequence-based prediction, we identified four out of eight predictions where SNPs inflicted damage to the structure. For structure-based prediction, we identified six out of six predictions where SNPs destabilized structures. We shortlisted ten mutations based on this consensus ranking approach. The outcomes of the sequence-based and structure-based predictions are presented in Supplementary Tables 3 and 4, respectively. Using these cut-offs, we identified and tabulated ten mutations for comparison. We further examined whether these nsSNPs (Table 3) elicited significant changes, consequently affecting the function of TG2.

**Table 3:** Analysis of nsSNP prediction. For each SNP the surface accessibility, location in the secondary structure, change in charged group, distance of the SNP location to other domains, distance of the SNP to the active site residues, and distance to the two calcium ions (labeled as I and II) are listed.

| S. No | Mutation | Initial structure (Relacc) | Final structure (Relacc) | Location of Secondary structure (DSSP) Initial | Location of Secondary structure (DSSP) Final | Change in charged groups | Distance b/w nsSNPs and Domain | Distance b/w nsSNPs and Active site | Distance b/w nsSNPs and Ca<sup>+2</sup> (I) | Distance blw nsSNPs and Ca<sup>+2</sup> (II) |
| --- | --- | --- | --- | --- | --- | --- | --- | --- | --- | --- |
| 1 | R48H | Buried | Intermediate | S - Bend | S - Bend | No Change | ~ 113.2 Å | ~ 63 Å | ~ 72.8 Å | ~ 54.6 Å |
| 2 | A110T | Buried | Buried | - | S - Bend | Change | ~ 80.1 Å | ~ 27.7 Å | ~ 42.8 Å | ~ 26.3 Å |
| 3 | E186Q | Buried | Exposed | S - Bend | S - Bend | Change | ~ 71.6 Å | ~ 19.1 Å | ~ 40.4 Å | ~ 25 Å |
| 4 | R214H | Buried | Buried | T - Turn | T - Turn | No Change | ~ 72 Å | ~ 23.4 Å | ~ 35.9 Å | ~ 26.1 Å |
| 5 | V224G | Buried | Buried | H - Helix | T - Turn | No Change | ~ 60.8 Å | ~ 10.1 Å | ~ 29.2 Å | ~ 25.6 Å |
| 6 | R271C | - | - | - | - | Change | ~ 56.6 Å | ~ 13.5 Å | ~ 29.5 Å | ~ 37.6 Å |
| 7 | C277S | Buried | Intermediate | H - Helix | H - Alpha helix | No Change | ~ 51 Å | 0 Å | ~ 29 Å | ~ 24.8 Å |
| 8 | E314K | Buried | Buried | H - Helix | H - Alpha helix | Change | ~ 25.3 Å | ~ 22.4 Å | ~ 26.6 Å | ~ 45 Å |
| 9 | I331N | - | - | - | - | Change | ~ 39.2 Å | ~ 9.5 Å | ~ 20.9 Å | ~ 32.2 Å |
| 10 | E549G | Buried | Exposed | H - Helix | G - 3 to 10 helix | Change | ~ 17.8 Å | ~ 37 Å | ~ 38.8 Å | ~ 63.2 Å |

Here, R48H nsSNPs have a large distance between the domain that shows flexibility and the residue location, whereas E549G nsSNPs have a smaller distance between the residue location and the domain showing flexibility. The changes in structure are shown in Table 5. Thus, based on the results obtained among 10 mutations, we hypothesize that the nsSNPs R48H, E186Q, C277S, and E549G are highly likely to cause a drastic change in TG2’s structure and its dynamics.

### Conclusion

Transglutaminase2 (TG2) protein has been known to be implicated in celiac disease (Klöck et al., 2011) Currently, two conformations of TG2 are available (open and closed) and there are reports of Ca^2+^ playing a role (Kim et al., 2020, Jang et al., 2014, Jeong et al., 2020, Lindemann et al, 2012). In this study, we explore the structural dynamics the human TG2 protein in the presence of Ca^2+^ ions. Currently, the specific transitions between the conformations are unknown.

In the molecular dynamics simulation, we explored four long time-scale setups: the open conformation with and without Ca^2+^, and the closed conformation with and without Ca^2+^. Each of these four setups were run for 200ns, within which we were able to observe domain-level conformational changes. Using an open conformation and closed conformation, we explored elastic network models to study the long-time scale dynamics of TG2. In the coarse-grained model of GNM, we observed inter and intra domain fluctuations that correlate with the ANM models. The motions of β1 and β2 domains were correlated in relation to other domains of TG2. Hotspot residues located near the active site in the open conformation. In the closed conformation, hotspot residues were located relatively far away from the active site residues. Relatively higher fluctuations were found in the core domain of TG2. Analyzing the molecular dynamics simulation trajectory, we observed large conformational changes in the open conformation without Ca^2+^, and few significant changes in the closed conformation without Ca^2+^. Our models describe the domain level conformational changes happening without Ca^2+^ ions. The presence of Ca^2+^ ions most likely restricts the domain movements of TG2. Among the available nsSNPs for TG2, we performed an exhaustive analysis of both sequence- and structure-based tools to estimate the effect of mutations on TG2. Using a consensus approach, we identified 10 nsSNPs that destabilize TG2 or are of deleterious in nature. This study sets the stage for further computational and experimental exploratory studies to understand the TG2 function and structural dynamics.

## Acknowledgements

DKS, KD, UV, and RMY acknowledge the SASTRA Deemed to be University for infrastructural support and research facilities. USDA is an equal opportunity provider and employer.

## Data availability

The data is made available in the manuscript using publicly available datasets.

## Author Contirubution Statement

D.K.S. and K. D. contributed to generating data, data analysis, and writing frist draft of the manuscript. T.Z.S contributed to ideation, conceptualization and overarching research goals. U.V. and R,M.Y also were involved in conceptualization and overarching research goals, supervision of the work, project administration and finalizing the manuscript.

## Supplementary Tables

**Supplementary Table 1:** List of closed conformation of TG2. The structures with missing residue information are listed for each.

| PDB ID | Ligand bound | Number of Missing Residues | List of Missing Residues |
| --- | --- | --- | --- |
| 6KZB | GDP | 6 | MET1 GLU84 - GLU85<br>LEU465 GLU469 - GLU470 |
| 6A8P | GTP | 14 | LYS464 - GLU470 GLU689 -<br>HIS695 |
| 1KV3 | GDP | 36 | MET1 - LEU14, HIS44 -<br>ASP55, THR123 - LEU132 |
| 4PYG | GTP | 10 | MET1 - GLU3, GLU689 -<br>HIS695 |
| 3LY6 | ATP | 14 | MET1 - GLU3, GLU84 -<br>GLU85, GLU689 - HIS697 |

**Supplementary Table 2:**
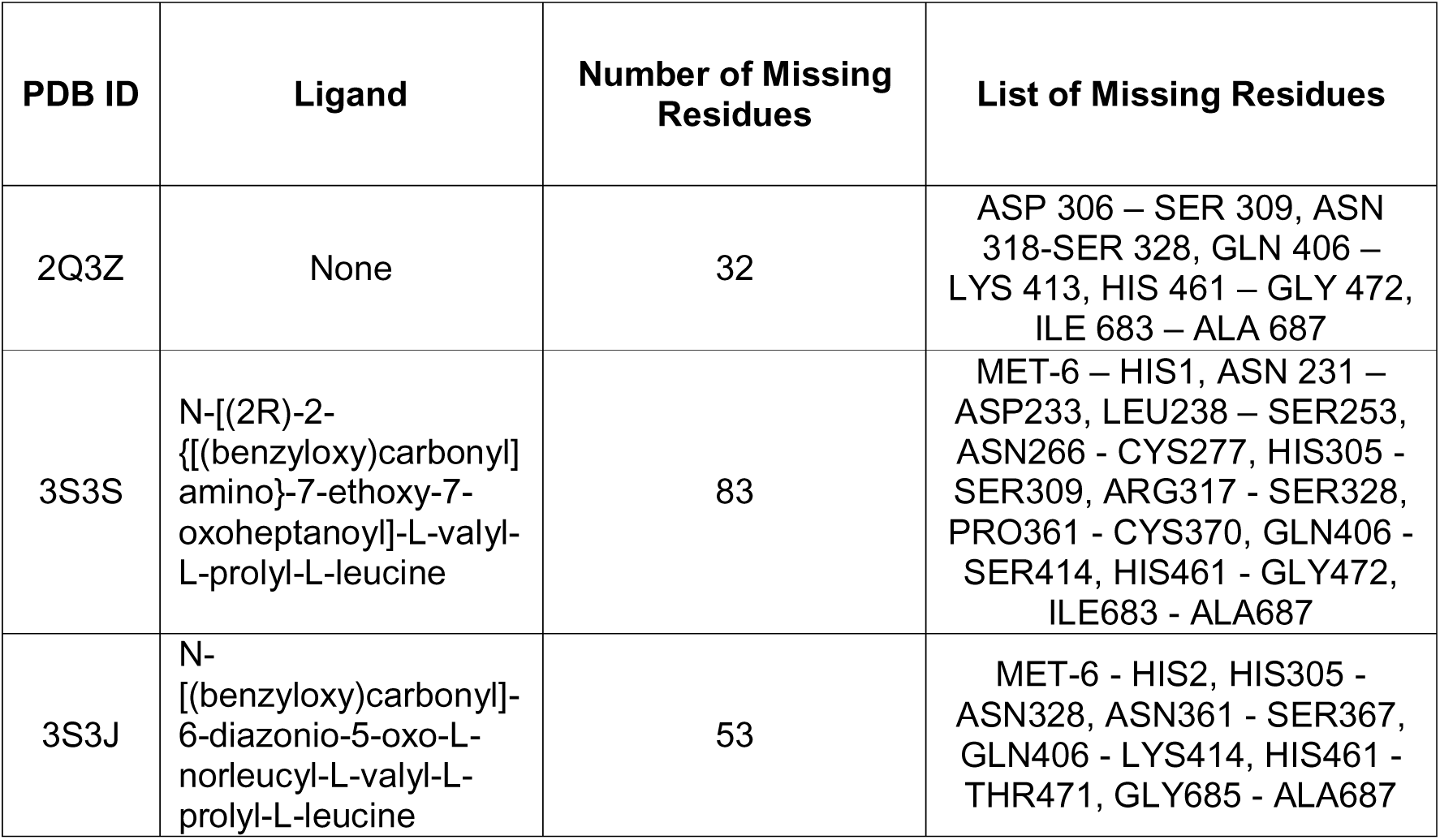
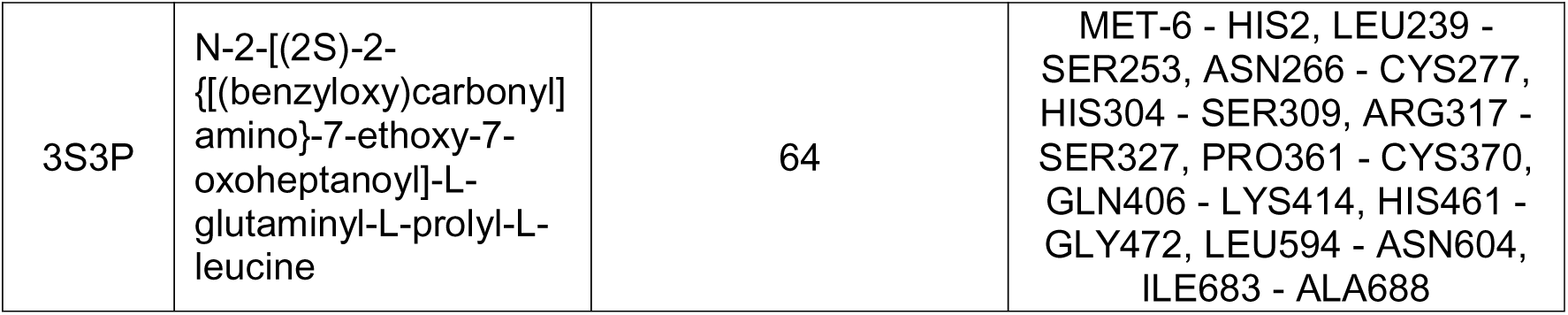
List of open conformation of TG2. The structures with missing residue information are listed for each.

**Supplementary Table 3:**
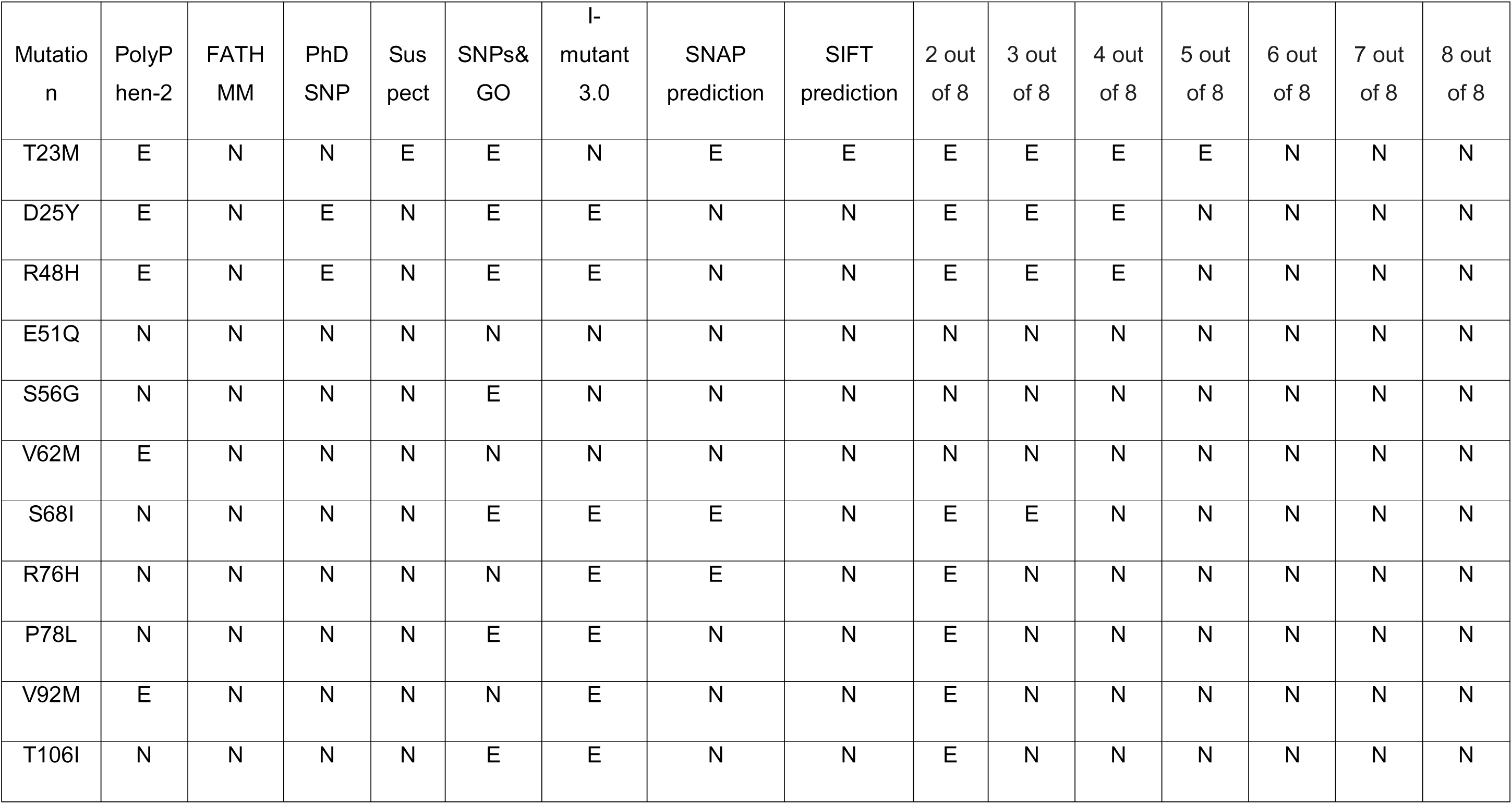
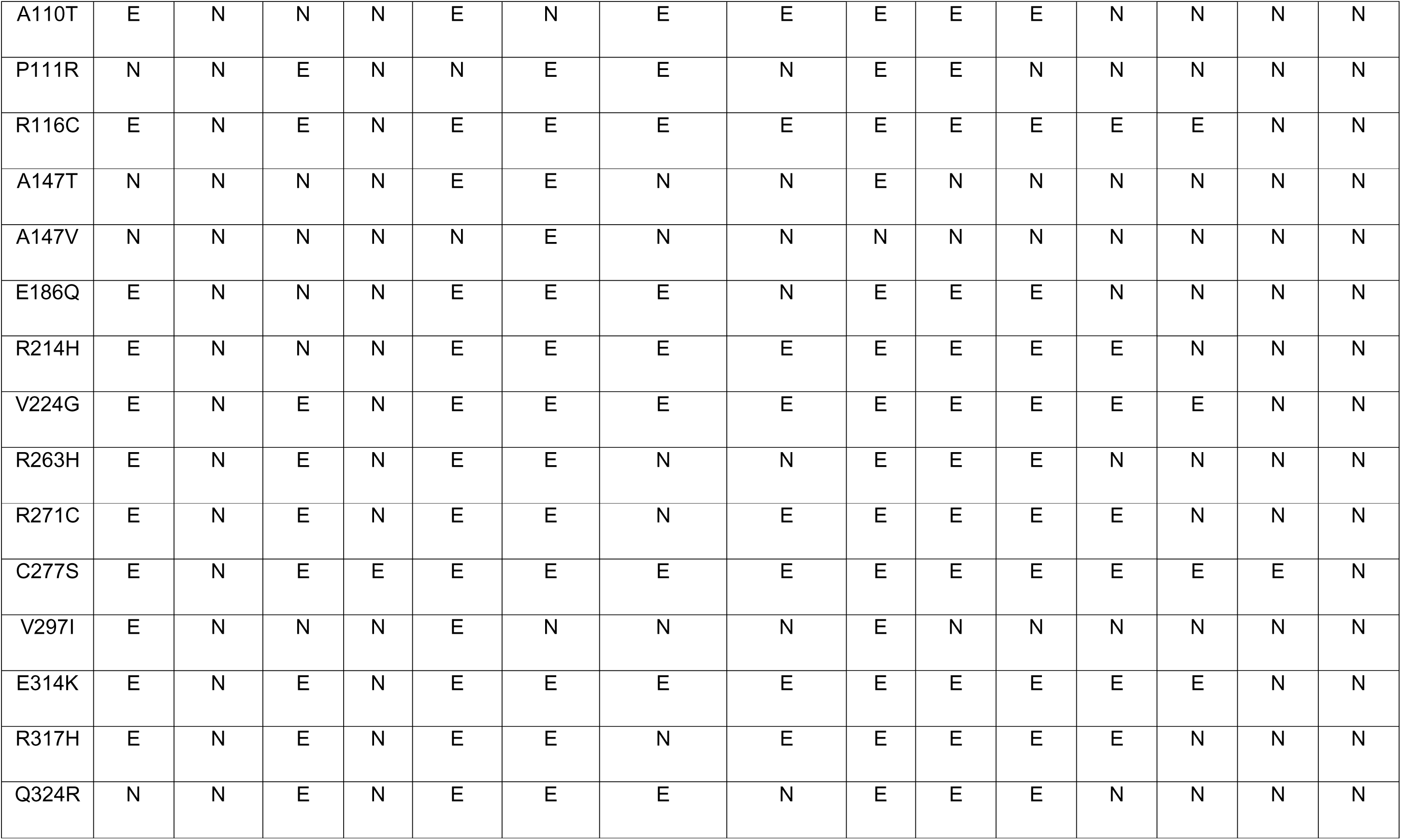
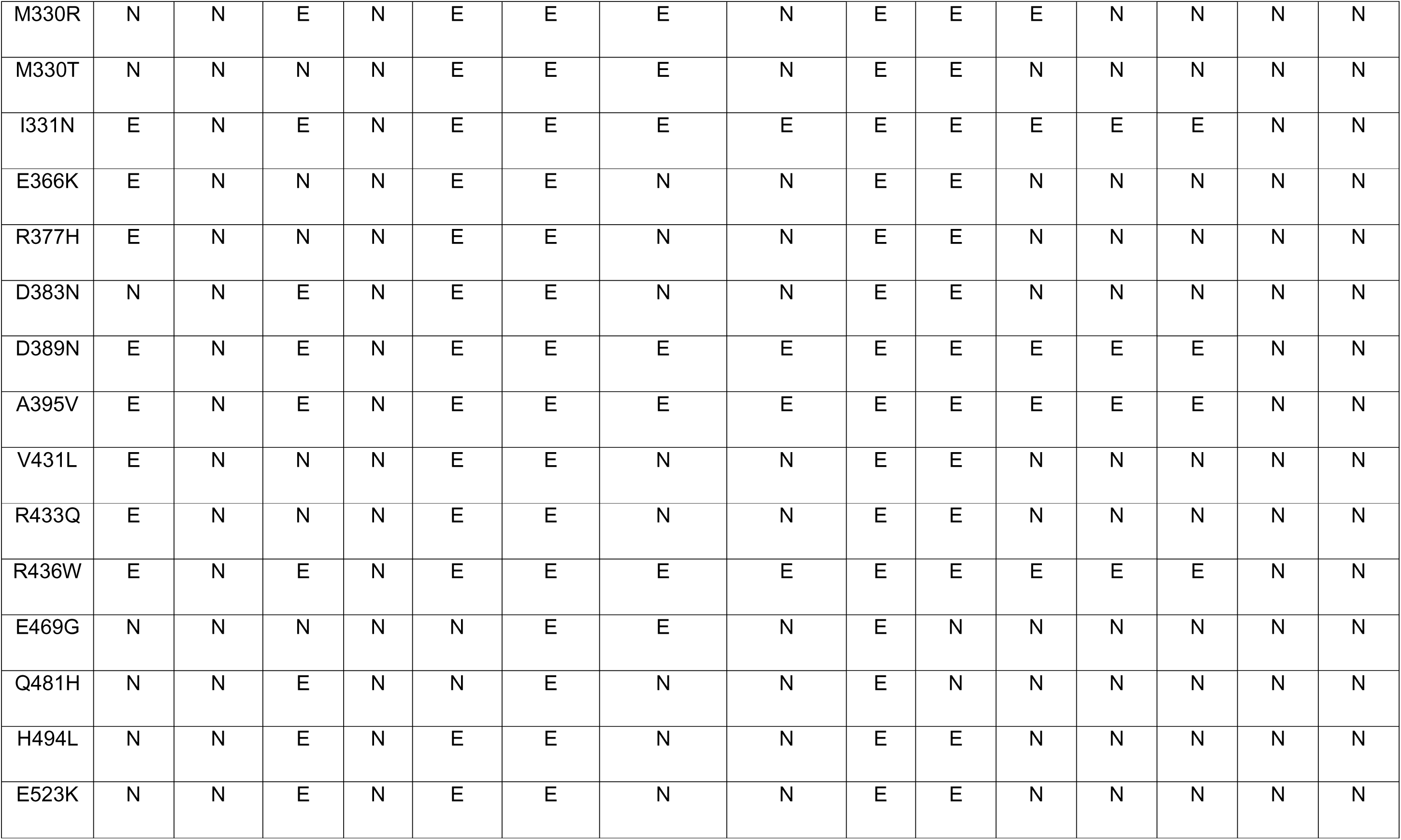
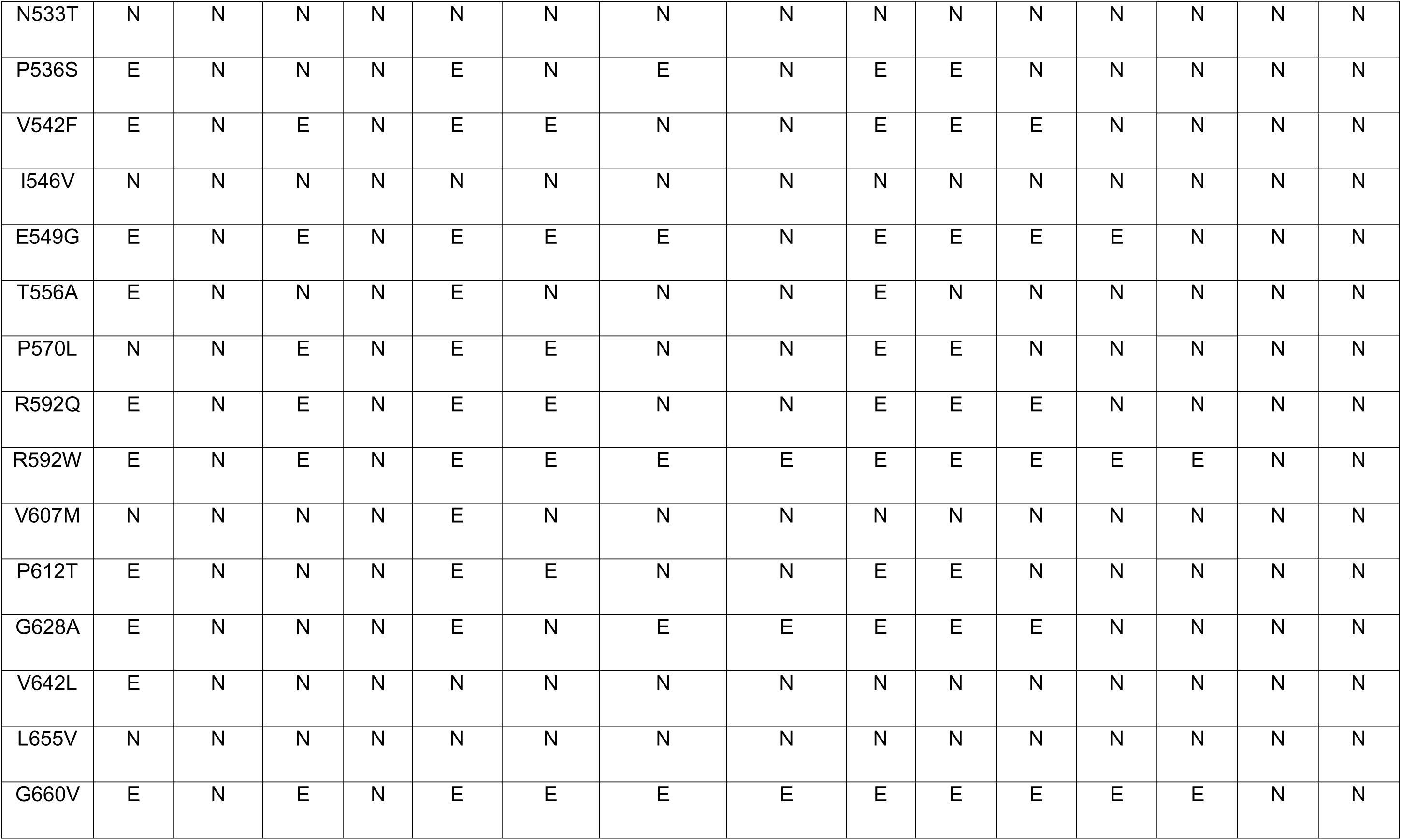
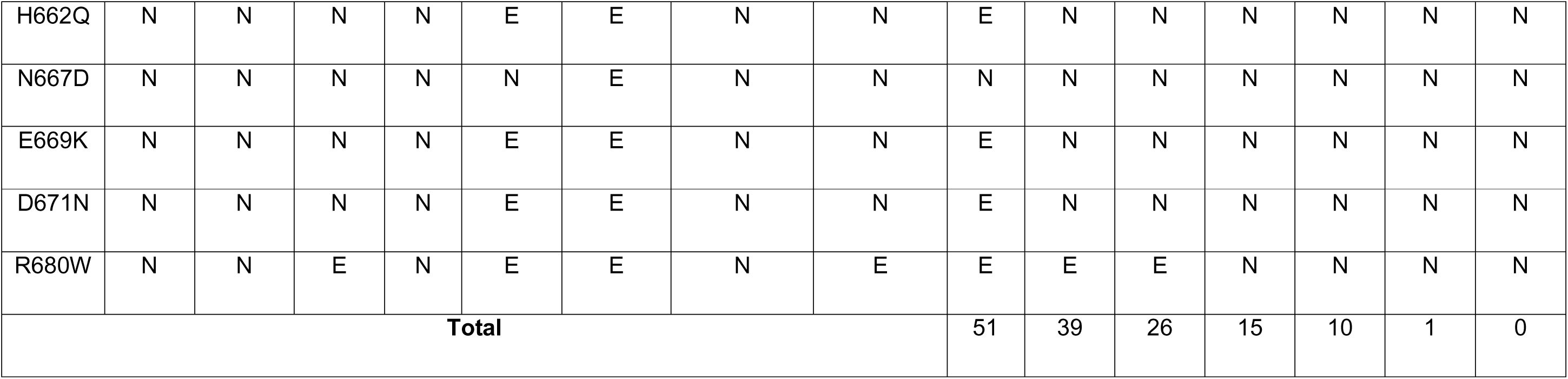
Sequence-based prediction of nsSNP by using various tools to check whether mutations have effect (E) or no effect (N).

**Supplementary Table 4 :**
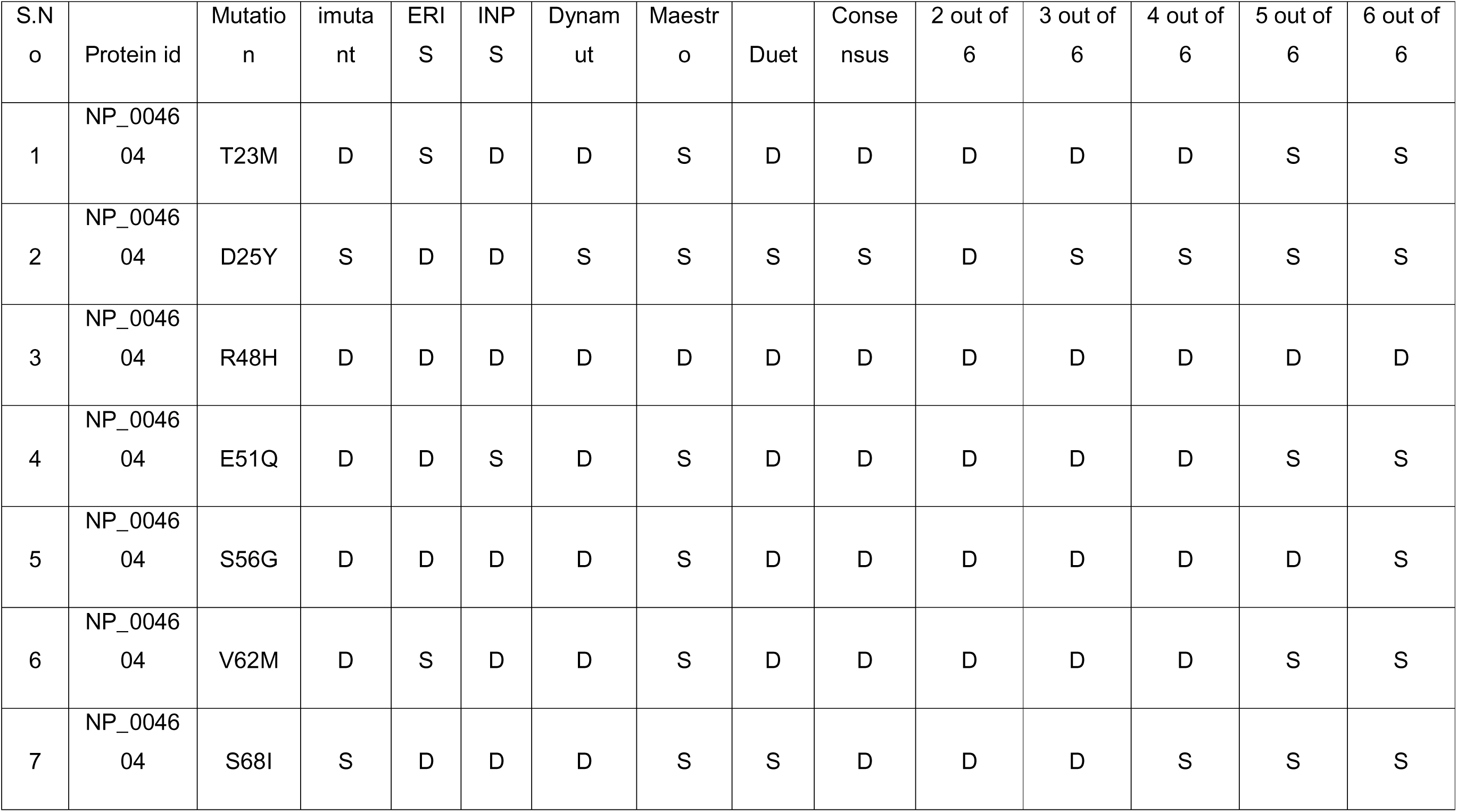
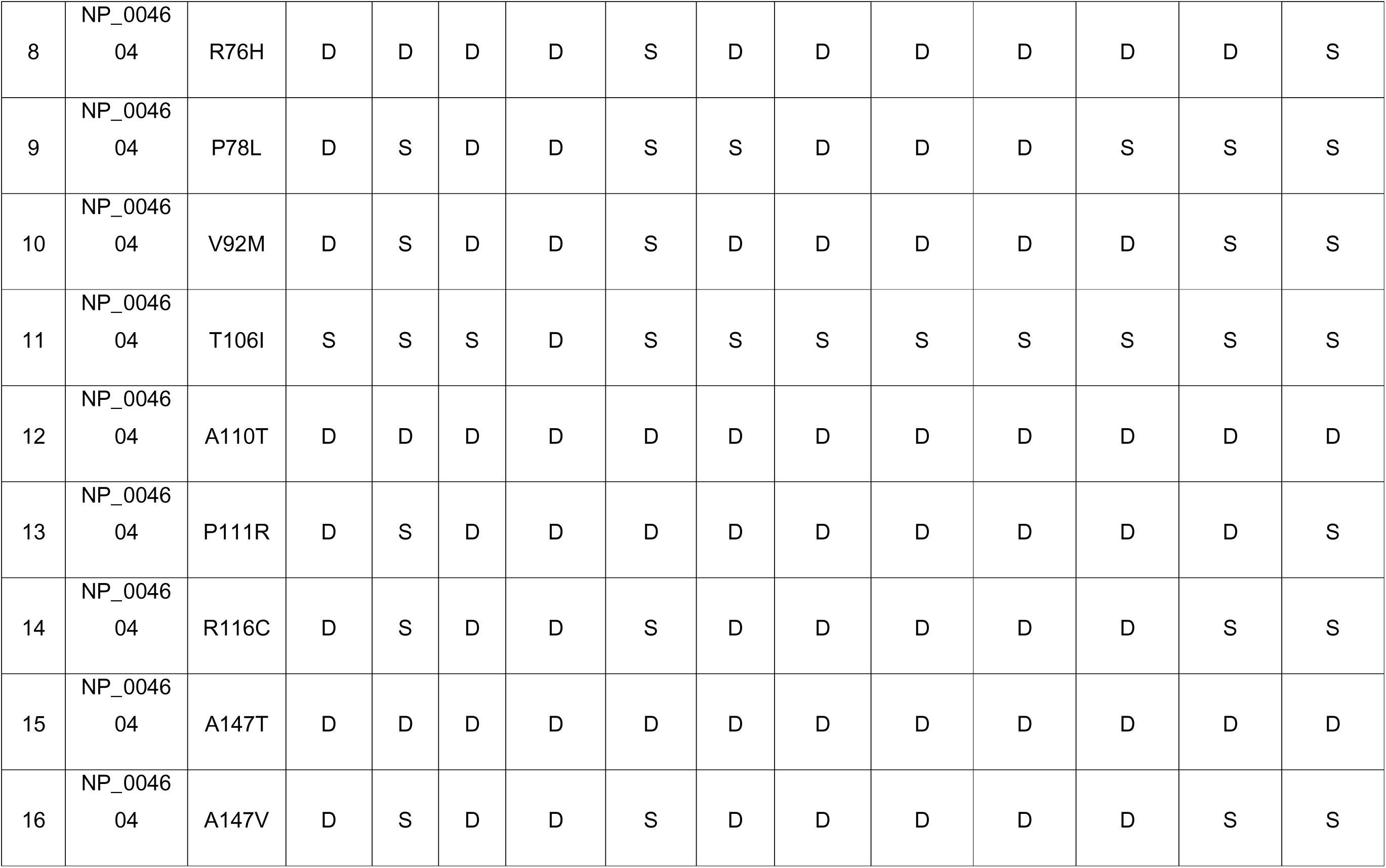
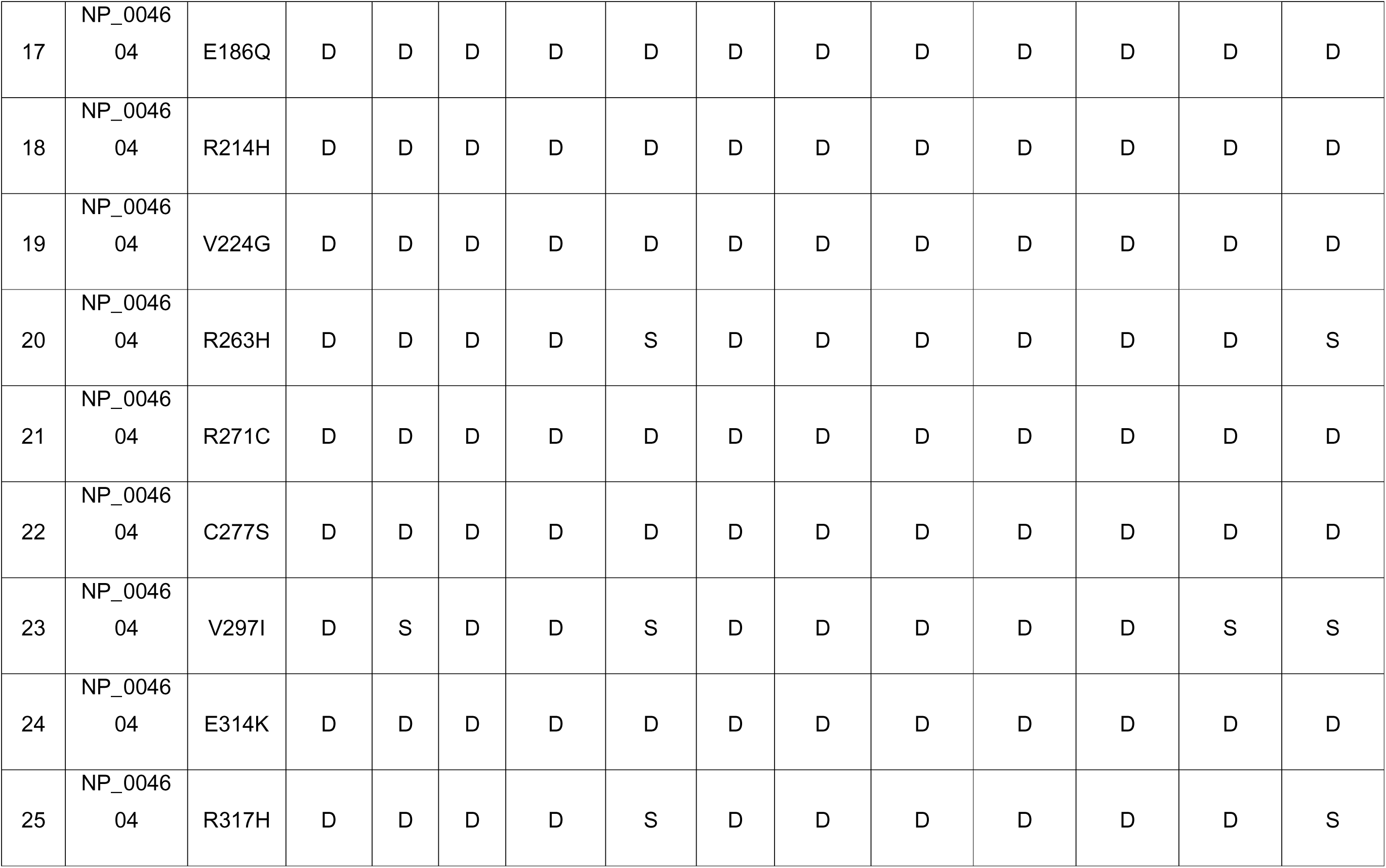
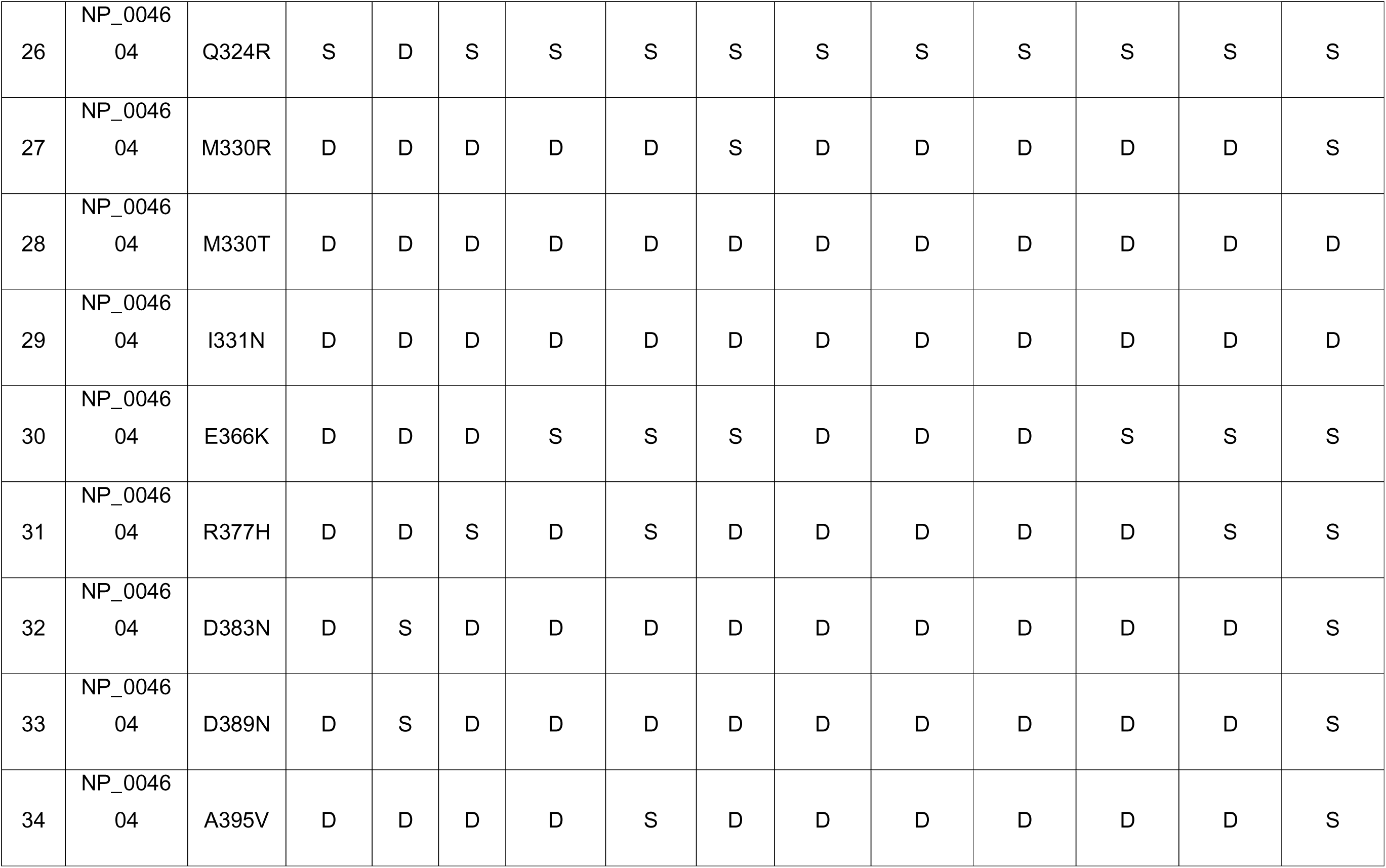
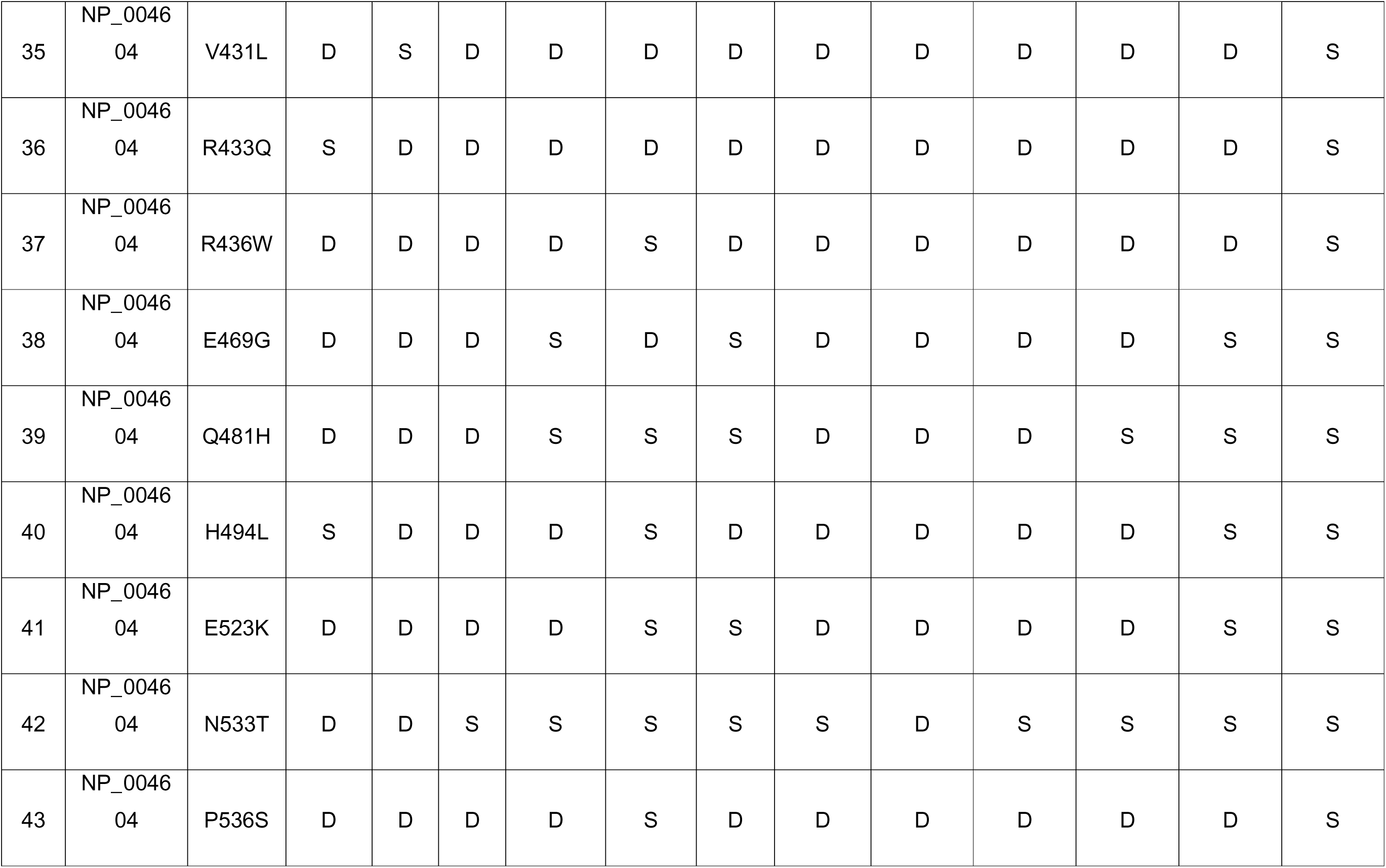
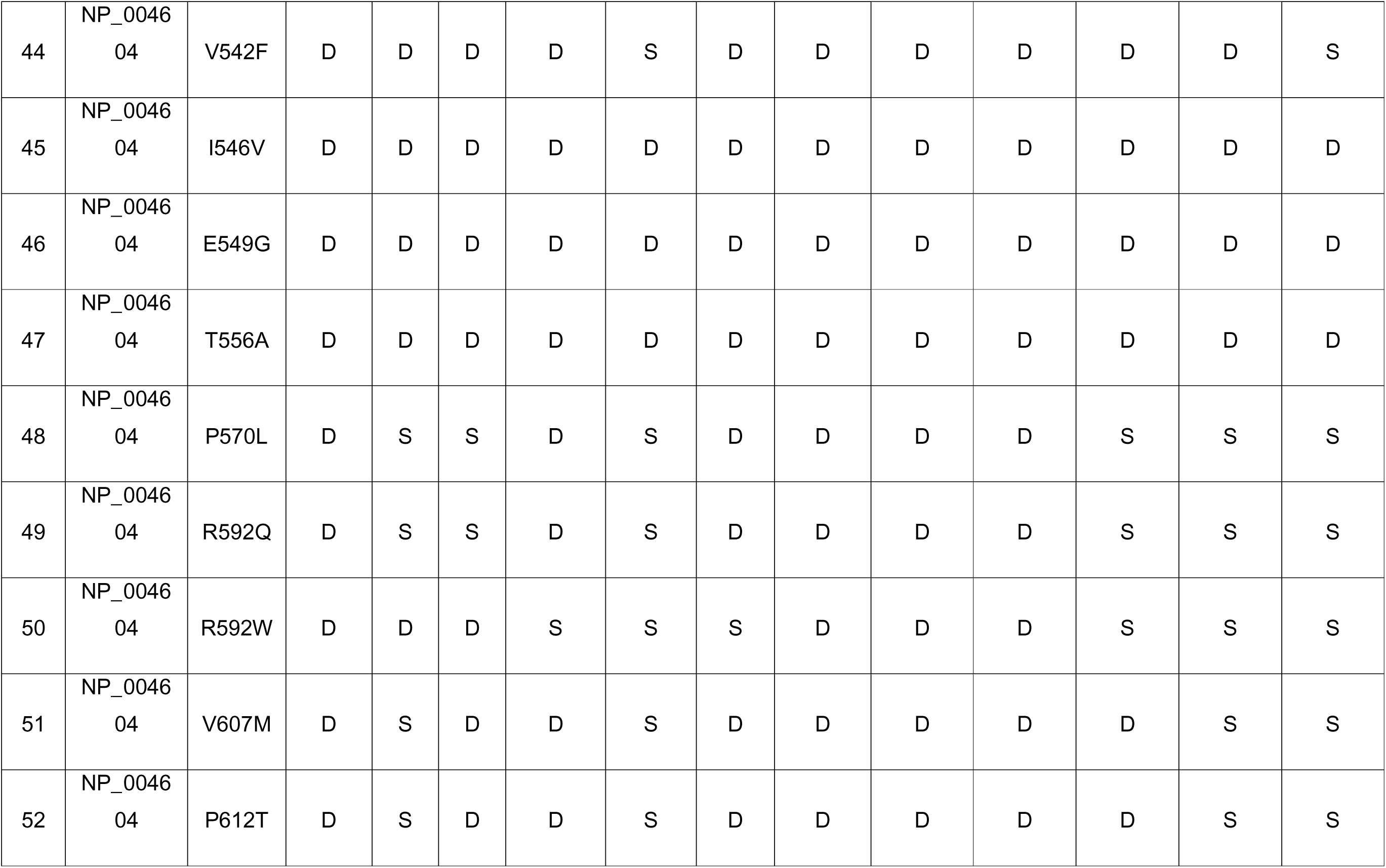
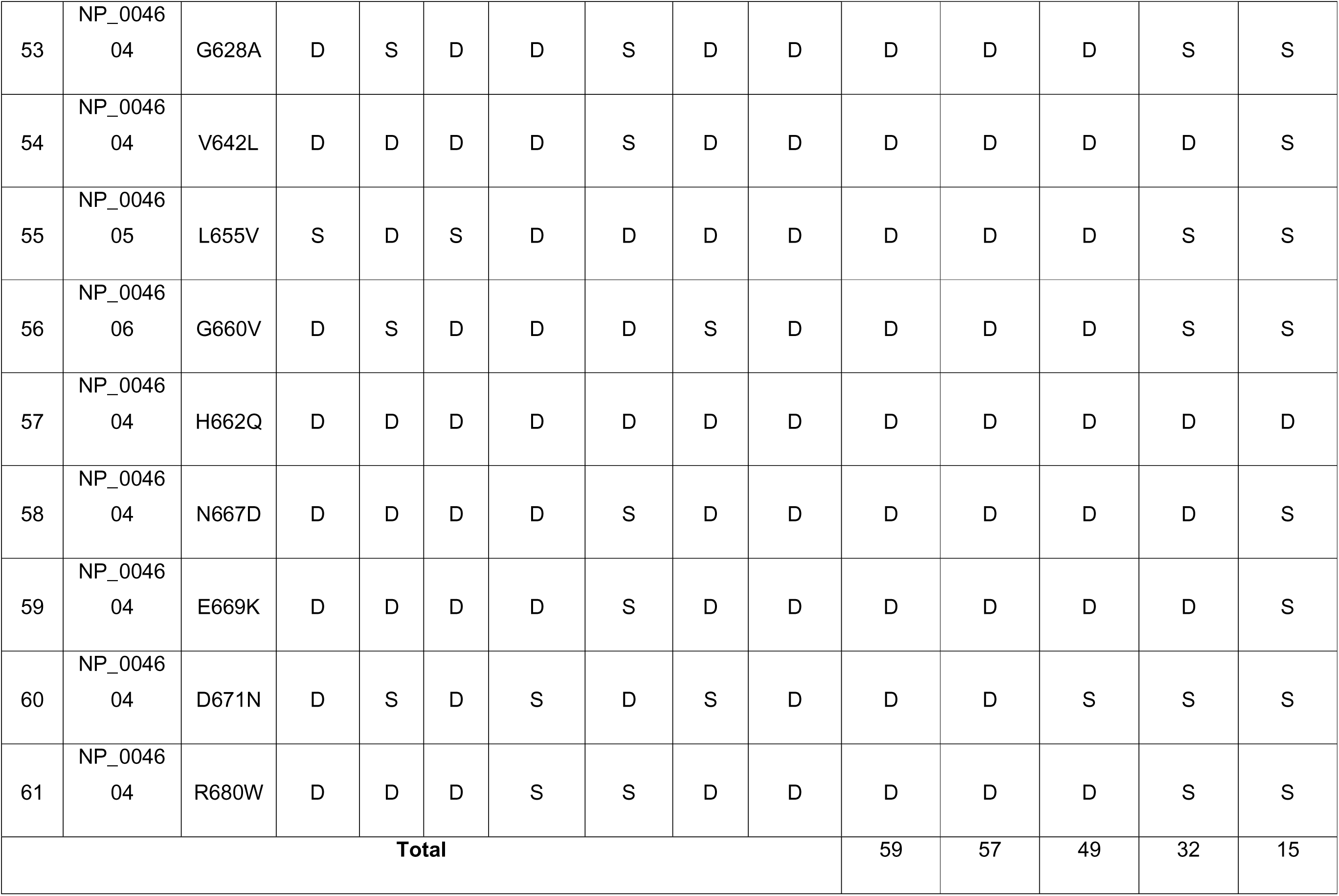
Structure-based prediction of nsSNP by using various tools to check whether mutations are stabilizing (S) or destabilizing (D).

## Supplementary Figures

**Supplementary Figure 1:**
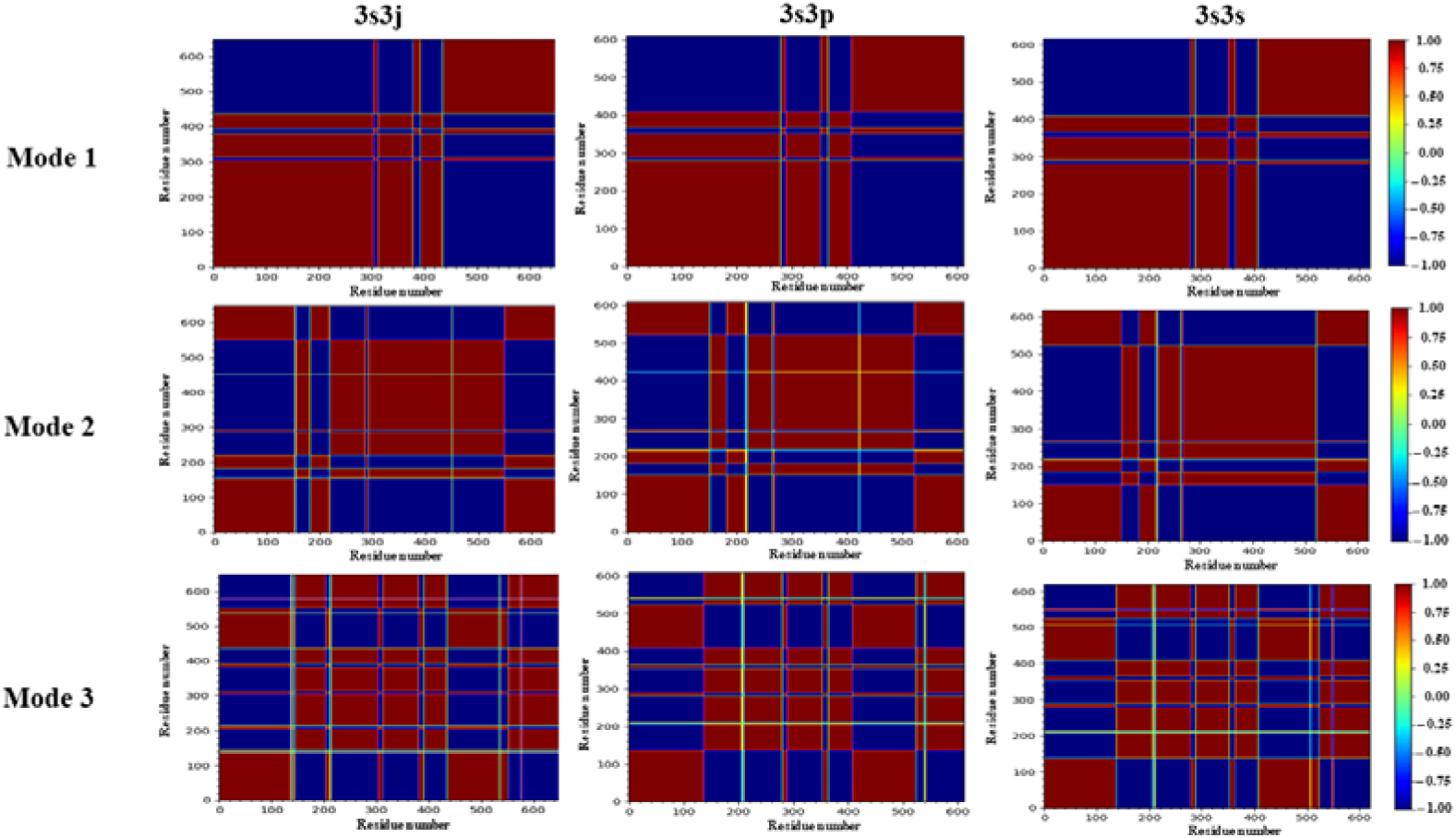
First three slowest mode of cross correlation plots from GNM (open conformation). The brown color block indicates the positively correlated and dark blue color block indicates the negatively correlated regions.

**Supplementary Figure 2:**
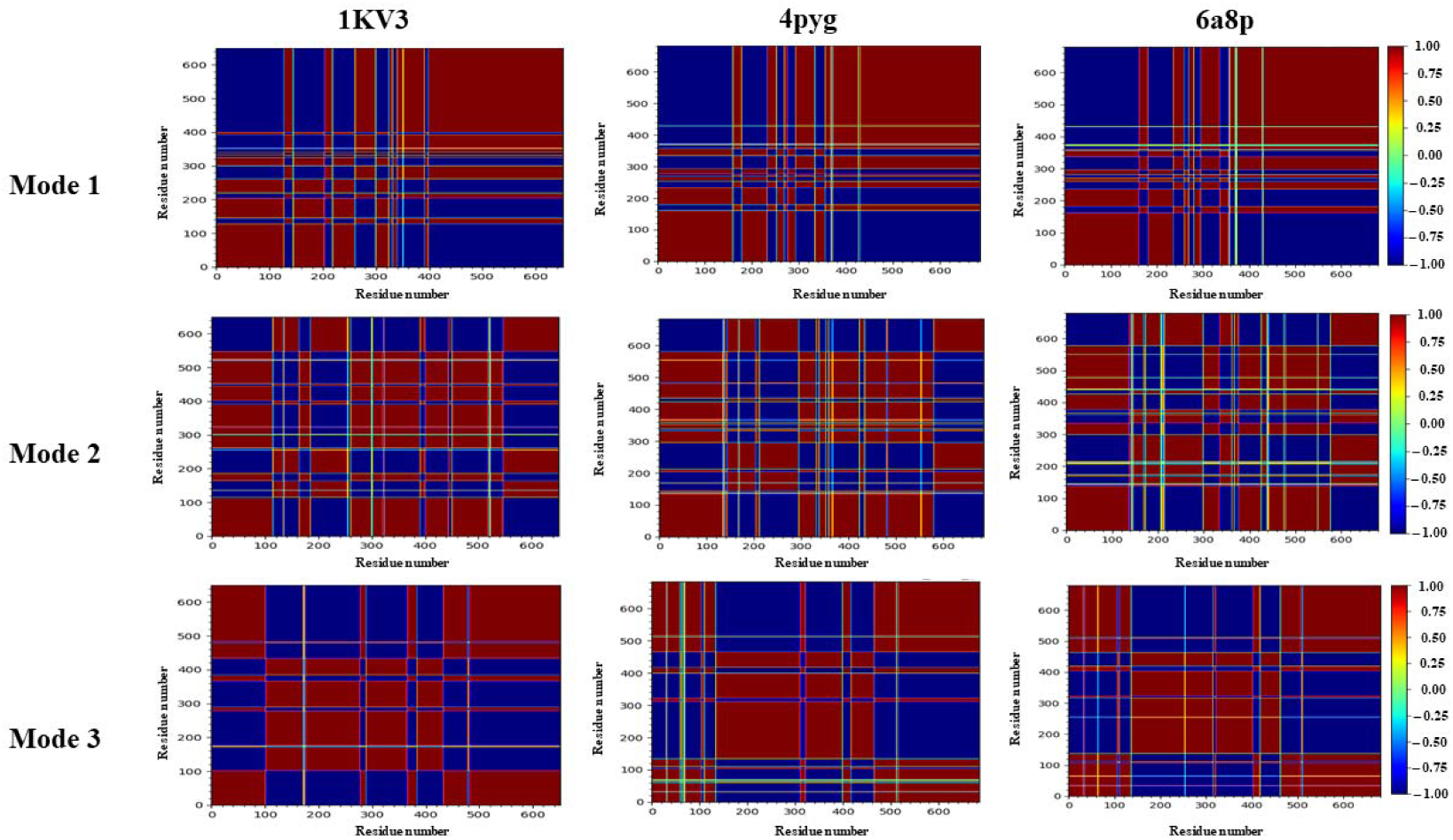
First three slowest mode of cross correlation plots from GNM (closed conformation). The brown color block indicates the positively correlated and the dark blue color block indicates the negatively correlated regions.

**Supplementary Figure 3:**
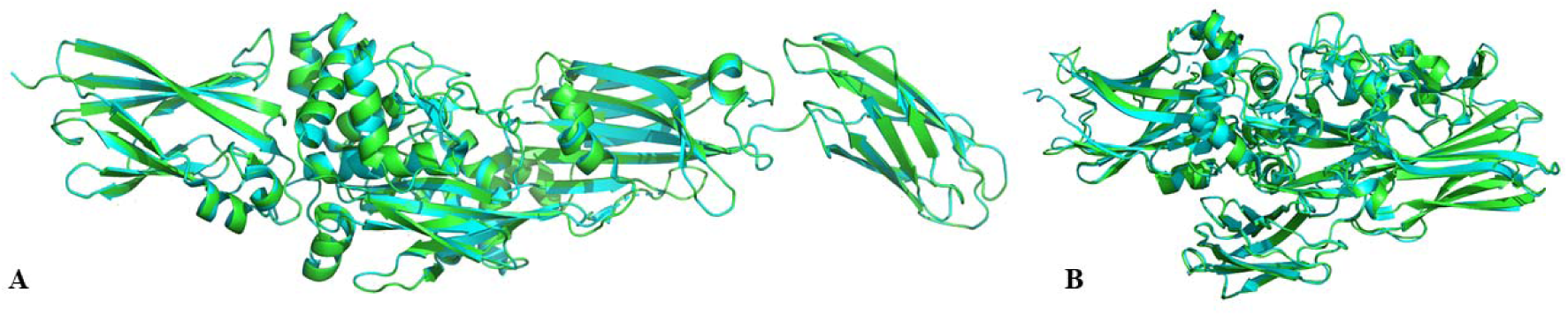
Superposing of (A) open and (B) closed conformation modelled structure with crystalline structure. Superposition made using align tool in PyMOL that gave an RMSD of 0.058 Å for the open conformation and 0.381 Å for the closed conformation. Here, green indicates crystal structure and cyan indicates modelled structures.

